# A potential anti-amyloidogenic therapy for type 2 diabetes based on the QBP1 peptide

**DOI:** 10.1101/2025.05.06.652241

**Authors:** María M. Tejero-Ojeda, Ada Bernaus Vives, Michal Wojciechowski, Dinh Quoc Huy Pham, Mateusz Chwastyk, Mario Vallejo, Anna Novials, Mariano Carrión-Vázquez

## Abstract

The self-assembly and aggregation of human islet amyloid polypeptide (hIAPP, or amylin) into β-sheet-rich structures, such as oligomers and fibrils, are implicated in pancreatic β-cell dysfunction and failure, contributing to the pathogenesis of Type 2 Diabetes (T2D). Consequently, extensive research has focused on identifying inhibitors, particularly short peptides, capable of targeting hIAPP and disrupting its amyloidogenic process, offering potential therapeutic strategies to prevent or slow T2D progression. In this study, we demonstrate the effectiveness of the anti-amyloidogenic peptide QBP1 in blocking the critical conformational transition to β-structure experienced by the hIAPP monomer, thereby in preventing amyloidogenesis and in reducing the cytotoxicity associated with its amyloid forms. First, we evaluated the anti-amyloidogenic effects of QBP1 through an *in vitro* aggregation methods, including a Thioflavin-T binding assay, dot blotting using the oligomer-specific A11 and fibril-specific OC antibodies, and negative staining electron microscopy. To assess its cytoprotective potential of QBP1 on hIAPP-induced toxicity, we examined its effects when fused to a protein transduction domain (penetratin) in INS-1E pancreatic β-cells, using viability assays and transcriptome analysis. Our results demonstrate that QBP1 effectively halts the formation of early toxic hIAPP intermediates, preventing amyloid progression and preserving β-cell viability and function. Additionally, molecular dynamics simulations revealed that QBP1 stabilizes amylin through strong van der Waals interactions and π-H bonds at hydrophobic and aromatic residues (*i.e.*, W, F), forming a stable binding network that prevents aggregation. Binding free energy analysis confirmed its high affinity, driven by favourable non-polar solvation energy and optimized structural complementarity. Collectively, these findings position QBP1 as a promising therapeutic candidate for preventing islet amyloid formation and mitigating β-cell dysfunction in T2D.

**Graphical Abstract:** 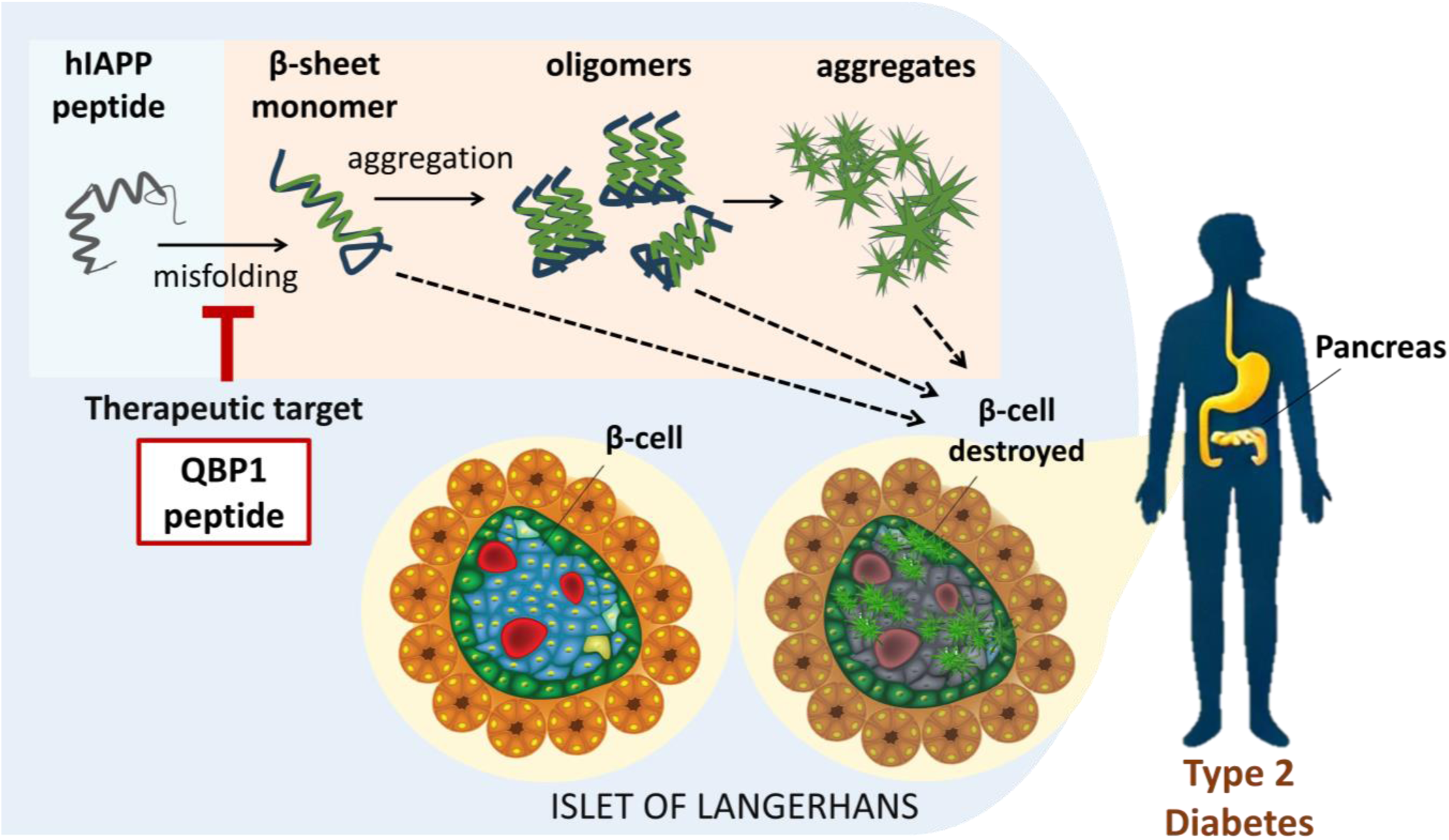

## Introduction

Amyloid diseases are a heterogeneous group of disorders characterized by the self-assembly and aggregation of proteins into toxic oligomers and insoluble fibrils, resulting in impaired physiological functions and tissue degeneration (Ross and Poirier, 2004). Among them, the aggregation-prone human islet amyloid polypeptide (hIAPP or amylin) is a major component of amyloid deposits in the pancreatic islets of patients with type 2 diabetes (T2D) (De Koning et al., 1994; Reed et al., 2021). This 37-residue peptide, co-secreted with insulin by pancreatic β-cells, contributes to glucose homeostasis by inhibiting insulin and glucagon secretion and modulating satiety (Akter et al., 2016; Ling et al., 2019). Although the hIAPP monomer is not inherently cytotoxic in humans, amyloid species formed during its pathological aggregation contribute to β-cell failure, depletion, and ultimately cell death, thereby accelerating the onset and progression of T2D (Cooper et al., 1988; Suárez et al., 2023). Soluble oligomers, intermediates in the aggregation pathways, have been identified as the most critical species (Kiriyama and Nochi, 2018), inducing cellular damage through membrane disruption and organelle stress (Back and Kaufman, 2012; Hernández et al., 2018). Furthermore, hIAPP hypersecretion has been associated with abnormal Ca²⁺ release and proteolytic enzyme activation, further exacerbating β-cell destruction (Alrouji et al., 2023; Kanatsuka et al., 2018).

The underlying molecular mechanisms responsible for hIAPP aggregation have been extensively studied (Alrouji et al., 2023; Clark and Nilsson, 2004), revealing that its amyloidogenesis is primarily driven by its central region, which contains hydrophobic amino acid residues crucial for their strong propensity to form β-sheet-rich structures (Bakou et al., 2017; Fortier et al., 2022; Wei et al., 2011). In contrast, rodent amylin possesses three proline residues—absent in the human protein—within this region, which enhance its solubility and prevent its pathological aggregation (Debbie et al., 2015; Raleigh et al., 2017). A key breakthrough in the development of therapeutic strategies was the substitution of positions 25, 28, and 29, by proline residues, which disrupt the β-structure of hIAPP. This modification led to the creation of Pramlintide (Symlin®), an FDA-approved hIAPP synthetic analogue used in the treatment of both type 1 and type 2 diabetes (T1D and T2D) (Boyle et al., 2022; McQueen and Bonk, 2005). Moreover, several “amyloidogenic hot spots” within the hIAPP sequence have been identified as promoters of cytotoxic β-sheet-rich folding intermediates (Mietlicki-Baase, 2018; Tenidis et al., 2000). Specifically, the β-sheet domains spanning residues 8–20, 20–29, and 30–37 have been implicated in fibril formation (Nilsson and Raleigh, 1999; Scrocchi et al., 2003) and are considered critical contributors to amyloidogenesis (Unnikrishnan and Shanmugam, 2022).

Given the pathological role of hIAPP amyloidogenesis, the development of inhibitors that effectively prevent hIAPP misfolding and subsequent amyloid formation without disrupting its physiological functions is of paramount importance. Short peptide inhibitors have emerged as a promising strategy in anti-amyloid therapy, offering high specificity, low toxicity, rapid clearance, efficient permeability, and excellent *in vivo* biocompatibility (Camus et al., 2008; Fernández Ramírez et al., 2023; Wang et al., 2023; Young et al., 2017). Several peptides, such as SNNFGA, GAILSS, NYGAILSS, and NFGAILPP, have demonstrated efficacy in blocking hIAPP aggregation *in vitro* by directly targeting its critical regions (Kapurniotu et al., 2002; Tenidis et al., 2000). Notably, the D-ANFLVH inhibitor has shown potential in preventing islet amyloid deposition, leading to reduced β-cell apoptosis and improved glucose tolerance in transgenic mouse models of T2D (Potter et al., 2009; Wijesekara et al., 2015). To further enhance efficacy, inhibitors incorporating breaker elements such as proline (Abedini et al., 2007) and α-aminoisobutyric acid (Aib) (Gilead and Gazit, 2004; Paul et al., 2017) have also been explored. Among these, proline substitution has proven particularly successful, as demonstrated by the aforementioned FDA-approved drug Symlin (Alrefai et al., 2010; McQueen and Bonk, 2005). However, despite its clinical effectiveness, Symlin faces challenges such as high production costs, limited solubility, and poor oral bioavailability (Alrefai et al., 2010). Newer amylinomimetics, such as Davalintide, show greater promise due to improved efficacy and longer-lasting effects, emphasizing the potential of short peptides in combating hIAPP amyloidogenesis (Boyle et al., 2022).

The PolyQ-binding peptide 1 (**QBP1**, core or minimal active sequence: **WKWWPGIF**) (Nagai et al., 2000; Ren et al., 2001; Tomita et al., 2009) was initially developed as an amyloidogenesis inhibitor targeting the early stages of polyglutamine (polyQ) aggregation (Popiel et al., 2013, 2011). QBP1 has been shown to interact with polyQ tracts in a random coil (RC) conformation, preventing their transition to β-sheet structures and thereby inhibiting oligomerization, amyloidogenesis, cytotoxicity, and neurodegeneration, as demonstrated in *in vitro* studies and in *Drosophila* and mouse models (Hervás et al., 2012; Nagai et al., 2003, 2007; Popiel et al., 2007; Yang et al., 2018). Although its systemic administration in a polyQ mouse model was previously hindered by proteolytic degradation and poor blood-brain barrier (BBB) penetration (Popiel et al., 2013, 2009; Takeuchi and Nagai, 2017), recent findings indicate that intranasal delivery significantly enhances its brain uptake and therapeutic efficacy (Takeuchi and Nagai, 2017). Beyond polyQ diseases, QBP1 has shown anti-amyloidogenic effects on other pathological amyloids, including α-synuclein (α-Syn) (Hervás et al., 2012) and TAR DNA-binding protein 43 (TDP-43) (Mompeán et al., 2019), as well as some functional amyloids such as the yeast prion domain Sup35 and the neuronal CPEB protein family (Hervás et al., 2021, 2016; Ramírez de Mingo et al., 2023). Interestingly, it must be noted that the aggregation of other amyloids, such as amyloid-β 42 (Aβ_42_) and tau protein, was not affected by QBP1 (Fernández Ramírez et al., 2023; Hervás et al., 2012).

Subsequent NMR studies have provided insights into the structural properties of QBP1, revealing that it adopts a preferred conformation consisting of a β-strand-like structure followed by a turn featuring a *trans* proline. This conformation is stabilized by a well-defined hydrophobic core formed by the side chains of tryptophan (W4) and isoleucine (I7), which is crucial for its inhibitory function (Ramos-Martín et al., 2014). Interestingly, a scrambled (SC) version of QBP1, referred to as **SC-M8** (**WPIWKGWF**) (Popiel et al., 2007), lacks both a preferred conformation and a defined hydrophobic core, and instead exists as a mixture of *cis* and *trans* isomers in the peptide bond preceding proline (Ramos-Martín et al., 2014). Despite these structural differences, SC-M8 may retain partial inhibitory capacity for some amyloids, suggesting that QBP1’s function is not solely dependent on its sequence and conformation but also on its amino acid composition, particularly its high W content (Mompeán et al., 2019; Tomita et al., 2009). The proposed mechanism of action for TDP-43 involves QBP1’s aromatic residues mediating hydrophobic interactions with amyloidogenic protein regions and disrupting intramolecular hydrogen bonds of glutamine- and asparagine-rich sequences. By interfering with these interactions, QBP1 prevents the stabilization and propagation of amyloid structures, highlighting its potential as a therapeutic agent for degenerative diseases like Amyotrophic Lateral Sclerosis (ALS) and Frontotemporal Dementia (FTD) (Fernández Ramírez et al., 2023; Mompeán et al., 2019).

In this study, we investigated whether inhibitory properties of the polyvalent QBP1 peptide could be extended to hIAPP, with the aim of blocking its critical conformational transition into a β-structure and preventing amyloidogenesis from the onset. To assess its effect on hIAPP aggregation, we performed time-resolved Thioflavin T (ThT) fluorescence spectroscopy, an amyloid-specific assay that detects fibril formation (Abedini et al., 2016). We also analyzed the morphology and abundance of hIAPP aggregates using complementary techniques, including immunodot blotting with anti-oligomer (A11) and anti-fibril (OC) antibodies (Kayed et al., 2007, 2003) as well as transmission electron microscopy (TEM). Furthermore, we assessed the ability of QBP1 to protect INS-1E pancreatic β-cells from hIAPP-induced cytotoxicity (Hohmeier et al., 2000; Soty et al., 2011) by measuring cell viability and the expression of key genes involved in β-cell function and stress responses (Casas et al., 2007; Rodríguez-Comas et al., 2020). To ensure efficient intracellular delivery, QBP1 was fused to the *Drosophila Antennapedia* (*Antp*, or penetratin) protein transduction domain (PTD), which allows receptor-independent membrane penetration (Popiel et al., 2009, 2007). Our results demonstrate that QBP1 effectively prevents the formation of early toxic hIAPP intermediates, halting amyloid progression and preserving β-cell viability and function. Molecular dynamics (MD) simulations provide additional insights into the underlying mechanism, showing that QBP1 stabilizes amylin through robust van der Waals forces and π-H interactions at critical hydrophobic and aromatic residues (such as tryptophan and phenylalanine). These interactions establish a stable binding network that hinders aggregation. Binding free energy analysis confirms its high affinity, driven by favorable non-polar solvation and structural complementarity. Overall, these results highlight QBP1 as a promising therapeutic agent for inhibiting islet amyloid formation and slowing the progression of T2D.

## Materials and Methods

### Peptide preparation

Synthetic peptides hIAPP (1–37), QBP1 (core sequence: WKWWPGIF) (Ren et al., 2001; Tomita et al., 2009), SC-M8 (WPIWKGWF) (Nagai et al., 2007), SC-M11 (WPIWSKGNDWF), a SC version of the original 11-residue QBP1 (Nagai et al., 2000), and *Antp*-QBP1 (RQIKIWFQNRRMKWKKGGWKWWPGIF) (Popiel et al., 2009) were synthesized by GenScript (Netherlands) at >95% purity. Following a disaggregation protocol, hIAPP peptide was dissolved in formic acid at a concentration of 0.5 mg/ml and incubated for 1 h at room temperature (RT) (Arnés et al., 2020). The solution was then subjected to 5 min of water bath sonication, aliquoted, dried under speed vacuum, and stored at −80 °C. Stock solutions of QBP1, SC-M8, SC-M11 and *Antp*-QBP1 peptides were prepared in DMSO at a final concentration of 10 mM and stored at −20°C until further use.

### Amyloid formation

Dried hIAPP peptide (0.5 mg) was reconstituted in 10 mM phosphate-buffered saline (PBS, pH 7.4) with 2% DMSO and 2 mM sodium azide to a final concentration of 100 μM. The solution was centrifuged at 16,000 x g for 50 min at 4 °C to remove any preformed aggregates, and the resulting supernatant (designated as “time zero”) served as the starting point for amyloidogenesis, consisting primarily of monomeric species. After vortexing, samples were incubated at 25 °C for 7 days without agitation to facilitate the gradual formation of oligomers and fibrils. QBP1 and its SC controls were added at varying molar ratios at the start of the incubation experiment to ensure their presence throughout the aggregation process. Aliquots were collected at different time points for biochemical characterization and biological assays in INS-1E β-cells. For cellular assays involving hIAPP oligomers, samples were sonicated prior to use to prevent incipient fibril formation (Kayed and Glabe, 2006; Patterson et al., 2019).

### ThT Fluorescence Assay

Amyloid formation was assessed *in vitro* using time-resolved fluorescence spectroscopy using ThT, following established protocols (Abedini et al., 2016; Gade Malmos et al., 2017). Reactions were carried out in PBS (pH 7.4), where hIAPP samples at varying concentrations were incubated with 25 µM ThT (prepared from a 1 mM stock solution). QBP1 and its SC controls were tested at five different molar ratios relative to hIAPP, while control samples (without additional peptides) contained equivalent DMSO concentrations. Fluorescence intensity (arbitrary units, a.u.) was monitored over 24 or 48 h at 30°C using a FLUOstar® Omega spectrophotometer (BMG LABTECH), with excitation and emission wavelengths set to 450 nm and 482 nm, respectively. Data were analyzed and fitted to non-linear models in OriginPro (OriginLab Corporation, Northampton, MA, USA) to compare aggregation dynamics. Key aggregation parameters, such as maximum fluorescence intensity (MFI) and half-time of aggregation (t₁_/_₂), were extracted to evaluate the extent and rate of amyloid formation (Gade Malmos et al., 2017).

### Fluorescence Microscopy

To further examine the structural organization of amyloid aggregates, fluorescence microscopy images of ThT-stained samples were acquired directly from the plate after 24 and 48 h using a Leitz DM IRB microscope (Leica, Germany) equipped with 10x/0.22, 20x/0.30 and 40x/0.8 objectives. For detailed imaging at 40x magnification, samples were then transferred to an Ibidi μ-Slide 4 Well ibiTreat chambered coverslip (Ibidi Gmbtt, Germany), which features a #1.5 polymer coverslip bottom that provides an optimal refractive index. At least three images *per* experimental condition were acquired and analyzed using Leica LAS AF Lite software.

### Confocal Microscopy and three-dimensional (3D) reconstruction of acquired images

For high-resolution visualization and 3D reconstruction of aggregates under different conditions, ThT-stained samples previously prepared in Ibidi μ-Slide 4 Well ibiTreat coverslips (Ibidi, Germany) were imaged using a Leica Stellaris 8 STED super-resolution microscope equipped with a 40x oil-immersion objective. For each experimental condition, at least two independent fields were imaged using laser excitation at [488 nm], with emission detected at [509-580 nm]. Z-stack acquisitions were performed to generate 3D reconstructions of aggregate structures. Image processing and quantitative analysis were conducted using ImageJ/Fiji (NIH, USA).

### Dot blot immunoassay

Two µL of incubated samples at various time points were spotted onto a nitrocellulose membrane. After blocking non-specific interactions for 1 h at RT with 10% non-fat milk (Blotting-Grade Blocker, Bio-Rad) in Tris-buffered saline containing 0.01% Tween 20 (TBS-T), the membrane was washed three times for 5 min with TBS-T and incubated for 1 h at RT with the A11 polyclonal anti-oligomer antibody (1:2000, ThermoScientific) (Kayed et al., 2003) and the fibril-specific OC polyclonal antibody (1:2500, Millipore) (Kayed et al., 2007) in 5% milk-TBS-T. After three 5-minute washes with TBS-T, the membrane was incubated for 1 h with the IRDye 680LT anti-rabbit secondary antibody (LI-COR) and then revealed in an Odyssey® CLx (LI-COR) imaging system. The final hIAPP concentration was 70 µM. Positive and negative controls consisted of 66 µM Aβ_42_ fibrils and 60 µM bovine serum albumin (BSA), respectively.

#### TEM

For ultrastructural analysis of the insoluble aggregates, 10 µL of incubated samples were adsorbed onto carbon-coated 300-mesh copper grids (Ted Pella Inc.) for 5 min, followed by negative staining with a 5% (w/v) aqueous uranyl acetate for another 5 min. The grids were washed twice with water and air-dried. Images were acquired at excitation voltages of 25 and 100 kV using a Jeol 1011 TEM equipped with a CCD Megaview III camera. Visualization of hIAPP amyloid fibrils was performed for a final concentration of 70 µM after 72 h of incubation. When present, inhibitors were tested at five different molar ratios relative to amylin.

### Viability Assays

Rat INS-1E pancreatic β-cell lines (INS-1E, INS-1E-hIAPP, and INS-1E-Ct) were used (Hernández et al., 2018; Hohmeier et al., 2000; Soty et al., 2011). The INS-1E-hIAPP cell line, which stably expresses human IAPP, and the INS-1E-Ct line, stably transfected with an empty vector, were used as experimental and negative controls, respectively. To assess the anti-amyloidogenic effect of QBP1 in these cell-based assays, we used the minimal active core of QBP1 (WKWWPGIF) fused to *Antp* PTD (RQIKIWFQNRRMKWKK) (Popiel et al., 2007), capable of penetrating cell membranes efficiently in a receptor-independent manner. Moreover, this peptide was biotinylated at the N-terminus for subsequent detection, and two central glycine residues were incorporated as flexible linkers to prevent steric hindrance and preserve structural integrity. The sequence used was then: Biotin-RQIKIWFQNRRMKWKKGGWKWWPGIF.

*-Maintenance of INS-1E β Cell Line.* Cells were maintained in RPMI 1640 medium (Thermo Fisher), supplemented with 10% heat-inactivated fetal bovine serum (FBS, Gibco), 11 mM glucose, 10 mM Hepes, 2 mM L-glutamine, 1 mM sodium pyruvate, 50 µM β-mercaptoethanol, and 1% penicillin-streptomycin (Thermo Fisher). INS-1E cells were incubated at 37°C in a humidified atmosphere containing 5% CO_2_. INS-1E-hIAPP and INS-1E-Ct cells were cultured under the same conditions, with the addition of 200 mg/ml Geneticin (Soty et al., 2011). The medium was changed every 2-3 days, and cells were passaged at approximately 80-90% confluence using trypsin-EDTA.

*-Cell Viability Under Overexpressed hIAPP Conditions.* INS-1E-hIAPP and control (Ct) cells were seeded at 200,000 cells/ml in 96-well plates and incubated for 16 h prior to treatment. After 48 h of exposure to different concentrations of *Antp*-QBP1 peptide, viability was assessed by measuring ATP levels using the CellTiter-Glo® assay (Promega), according to the manufacturer’s instructions. Briefly, the plates were equilibrated at RT for 30 min, and the CellTiter-Glo® reagent was mixed with the cell supernatant at a 1:1 ratio. After a 10-minute incubation at RT with shaking, luminescence was measured using a FLUOstar® Omega microplate reader (BMG LABTECH). Wells containing only medium without cells were included to determine background luminescence. Additionally, phase-contrast microscopy was used to monitor morphological changes in cells following *Antp*-QBP1 treatment. Images were acquired using a Leitz DM IRB microscope (Leica, Germany) equipped with a Peltier temperature controller (Linkam, UK) set to 37°C, and a Leica K5 camera (Leica, Germany). At least three fields *per* condition were acquired using 10x/0.22 and 20x/0.30 objectives and analyzed with Leica LAS AF Lite software.

*-Cell Viability Under Extracellular hIAPP Oligomer Exposure.* INS-1E cells were cultured in high-glucose medium (16.7 mM) (Back and Kaufman, 2012; Lin et al., 2019) to simulate T2D-associated hyperglycemia and seeded at 200,000 cells/ml in 96-well plates, followed by 16 h of incubation. After 48 h of exposure to hIAPP prefibrillar oligomers (A11-positive) or buffered controls, with or without *Antp*-QBP1, cell viability was assessed using the CellTiter-Glo® assay. For gene expression analysis, the same treatments were applied in P6 Corning plates to maintain experimental consistency.

### Immunocytochemistry (ICC)

ICC was performed to evaluate hIAPP overexpression in INS-1E-hIAPP cells, confirm its absence in controls, and verify the cellular uptake of *Antp*-QBP1 following exogenous administration. Cells were seeded in 24-well plates on 12 mm coverslips pre-treated with 1 mg/ml poly-D-lysine (PDL) in borate buffer (pH 8.0) at a density of 200,000 cells/ml. After 48 h of incubation in culture medium containing either 10 µM biotin-tagged *Antp*-QBP1 or a control solution (Popiel et al., 2007), cells were fixed with 4% paraformaldehyde (PFA) for 15 min at RT. Then, cells were permeabilized with 0.1 % Triton X-100 in PBS for 15 min and incubated with blocking solution (2 % BSA, 5 % normal goat serum (NGS), 0.1 % Triton-X100 in PBS) for 1 h at RT. Cells were then incubated overnight at 4°C with primary antibodies against hIAPP (rabbit anti-hIAPP, 1:400; Sigma) and biotin (mouse anti-biotin, 1:200; VECTOR). Fluorophore-conjugated secondary antibodies were applied for 1 h at RT. Coverslips were mounted on slides using ProLong® Gold mounting medium (Life Technologies), and images were captured using a LEICA DMI 6000 fluorescence microscope equipped with a 40x/1.00 oil-immersion objective. Image analysis was conducted with ImageJ/Fiji software (NIH, USA), with a minimum of three images analyzed *per* experimental condition.

#### Gene expression analyses

*-RNA isolation.* Total cellular RNA from hIAPP-treated cells was isolated using the NzyTech Total RNA Tissue Kit (NZYtech) according to the manufacturer’s protocol, including on-column DNase digestion with the RNase-Free DNase Set (Promega). RNA quality and concentration were assessed using a Nanodrop One spectrophotometer (Thermo Scientific). Three independent experiments were performed to ensure reproducibility.

*-Real-time quantitative PCR (RQ-PCR).* One µg of total RNA was reverse-transcribed into complementary DNA (cDNA) using the High-Capacity cDNA Reverse Transcription Kit (Thermo Fisher). RQ-PCR was conducted with 0.5 ng of cDNA per reaction using SYBR Green Reagents (NZYtech) and a QuantStudio™ 3 RQ-PCR System (ThermoFisher), following the manufacturer’s protocol. The oligonucleotide primers used for gene expression analysis of *Pdx1*, *IL-1β*, *Slc2a2*, *Hspa5*, and 18S rRNA (used as the reference housekeeping gene) were obtained from Merck KGaA (Darmstadt, Germany) and are listed in **Table S1, *Supplementary material***. Gene expression levels were analyzed using the ΔΔCt method, normalizing to 18S rRNA expression.

### Statistical Analysis

All data are presented as mean ± standard error of the mean (SEM) from at least three independent experiments. Statistical comparisons were performed using one-way or two-way analysis of variance (ANOVA), as appropriate, followed by Bonferroni’s post-hoc test for multiple comparisons. A significance threshold of *p <* 0.05 was applied for all analyses. Normality and homogeneity of variance were verified prior to statistical testing. All statistical analyses and graphical representations were carried out using GraphPad Prism software (version 8.00; GraphPad Software, La Jolla, CA, USA) and OriginPro (OriginLab Corporation, Northampton, MA, USA).

### MD simulations

To investigate the binding interactions between hIAPP, a 37-amino-acid peptide with a disulfide bridge (CYS2–CYS7), and QBP1 (WKWWPGIF) (Ramos-Martín et al., 2014), together with its SC variants, SC-M11 (WPIWSKGNDWF) and SC-M8 (WPIWKGWF), molecular docking and MD simulation studies were conducted. The amylin structure under SDS micelles at pH 7.3, was obtained from the Protein Data Bank (**PDB: 2L86,** depicted in **Figure S1, *Supplementary material***) (Nanga et al., 2011) and served as the starting point for the simulations. Complexes were generated *via* molecular docking using the HADDOCK server (De Vries et al., 2010), with peptides capped at both termini (N-terminal acetyl and C-terminal amide). The docking process yielded 10 of the most probable configurations, which were subsequently used as initial structures for simulations to generate snapshot ensembles for binding analysis. To ensure reliable results, each configuration was simulated for 100 nanoseconds (ns) across 10 trajectories, with conformations saved every 10 picoseconds (ps), resulting in a total of 10,000 structures for analysis.

Simulations were performed using the Amber20 package (Case et al., 2020) with the AMBER 19SB force field (Tian et al., 2020) and the explicit solvent OPC water model (Izadi et al., 2014). The docked structures were positioned at the center of a simulation box with a buffer of at least 10 angstroms (Å) from the box edges to prevent interactions with periodic images, applying a 9 Å cutoff for non-bonded interactions. To mimic physiological conditions, the system was neutralized by adding Na⁺ and Cl⁻ ions, and the ionic strength was adjusted to 0.15 M. The simulations were conducted using the leap-frog integration algorithm (Leimkuhler and Matthews, 2013) with a time step of 2 femtoseconds (fs), while the SHAKE algorithm (Ryckaert et al., 1977) was applied to constrain hydrogen bonds, enhancing computational efficiency. Temperature was maintained at 300 K using a Langevin thermostat (Brooks et al., 1983) with a collision frequency of 2 ps⁻¹ to regulate kinetic energy and ensure system stability. Long-range electrostatics interaction were calculated using the particle mesh Ewald (PME) method (Darden et al., 1993).

Before the production run, the system underwent three preparatory steps: (1) energy minimization of the solvated amylin-QBP1 complex to eliminate steric clashes and optimize the initial structure, (2) gradual heating to 300 K under the NVT ensemble (constant number of particles, volume, and temperature) for controlled temperature stabilization, and (3) equilibration under the NPT ensemble (constant number of particles, pressure, and temperature) to reach a target density of approximately 1.0 g/cm³. Following these steps, conformations for analysis (Roe and Cheatham, 2013) were collected after 500 ps of equilibration in the NVT ensemble to ensure structural stability. For data analysis, contact maps (Gunnoo et al., 2018) were generated following the overlap criteria described by Mioduszewski et al. (2023) (Chwastyk et al., 2015; Mioduszewski et al., 2023), while binding energies were computed using the MM/PBSA (Molecular Mechanics Poisson-Boltzmann Surface Area) approach (Case et al., 2016), which estimates the free energy of binding by considering both enthalpic and entropic contributions.

## Results

### QBP1 strongly inhibits hIAPP amyloid fibril formation from the start of the process

To evaluate the *in vitro* amyloidogenic potential of hIAPP and the inhibitory effect of QBP1 on it, ThT-based fluorescence assays were performed for 24 h. As shown in **Figure 1A**, the aggregation kinetics of hIAPP exhibited a sigmoidal profile characteristic of amyloid polymerization, consisting of three distinct phases: a short lag phase, a rapid growth phase, and a steady-state phase in which the soluble peptide reached equilibrium with the fibrils. The aggregation process displayed a concentration-dependent acceleration, where higher peptide concentrations (50 μM, 70 μM, and 100 μM) resulted in a shorter lag phase and an increased fluorescence signal at the steady-state phase, indicating enhanced amyloid nucleation and fibril formation. The buffer control showed no detectable ThT fluorescence, confirming the specificity of the signal for amyloid aggregation.

**Figure 1.**
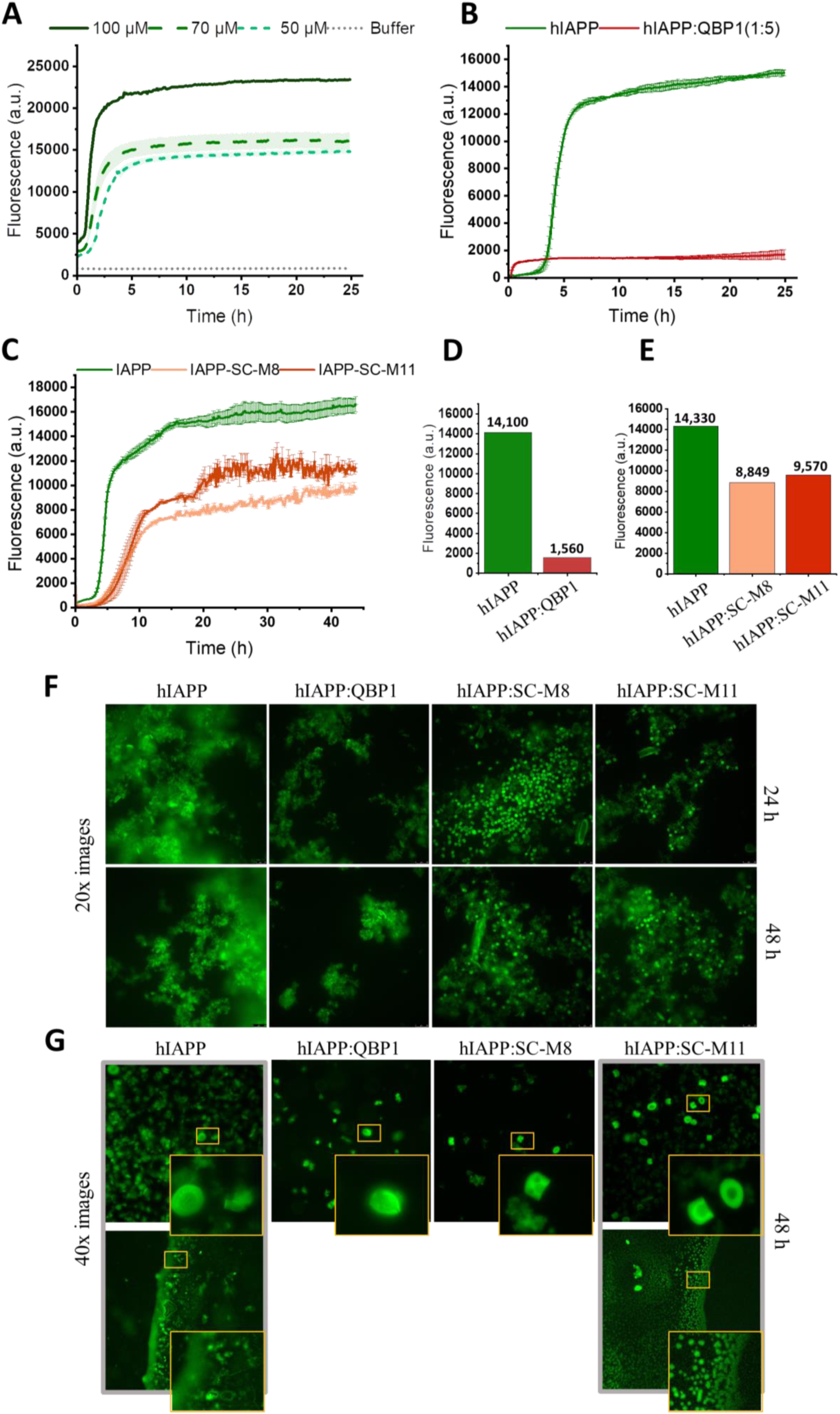
Inhibition of early-stage hIAPP amyloid fibril formation by QBP1 and its scrambled variants. **(A)** ThT fluorescence kinetics of hIAPP at increasing concentrations (50, 70, and 100 μM) over 24 h. Higher peptide concentrations accelerated both the lag and elongation phases compared to the buffer control, confirming concentration-dependent amyloidogenesis. **(B)** Aggregation kinetics of hIAPP in the presence of QBP1 (1:5 molar ratio). ThT fluorescence was drastically reduced, indicating strong inhibition of fibril formation by QBP1. **(C)** Effect of SC-M8 and SC-M11 (1:5 molar ratio) on hIAPP aggregation. Both SC variants decreased fluorescence, albeit less effectively than QBP1. **(D-E)** Maximum fluorescence intensity (MFI) values extracted from (B) and (C), confirming the inhibitory capacity of QBP1 and partial inhibition by SC variants. **(F)** Fluorescence microscopy images (20×) of ThT-stained samples after 24 and 48 h. hIAPP alone formed dense, clustered ThT-positive aggregates; QBP1-treated samples showed markedly reduced fluorescence, while SC-treated conditions exhibited intermediate signal intensity and aggregate abundance. **(G)** Higher-magnification images (40×) at 48 h reveal distinct aggregate morphologies. Small spherical structures were observed in both hIAPP and hIAPP: QBP1 conditions (*top insets, first and second columns*), potentially corresponding to immature or off-pathway intermediates. QBP1 reduced aggregate size and organization, while SC peptides promoted the formation of well-defined spherical or ring-like structures (*top insets, third and fourth columns*). Notably, SC-M11 samples exhibited droplet-like condensates at the drying interface (*bottom inset, fourth column*), indicative of phase separation. Insets show higher-magnification views of selected aggregates. Representative data and images from three independent experiments.

In contrast, the presence of QBP1 at a 1:5 molar ratio (hIAPP: QBP1) drastically reduced the fluorescence signal from the onset, indicating a strong inhibition of amyloid polymerization **(Figure 1B)**. To quantify this effect, the data were fitted to non-linear models in OriginPro, allowing for a detailed comparison of aggregation dynamics (Gade Malmos et al., 2017). The fluorescence intensity profile of <u>hIAPP alone</u> followed a <u>Boltzmann model</u>, whereas that of hIAPP in the presence of <u>QBP1</u> was fitted to a <u>Michaelis-Menten model</u>, suggesting distinct kinetic behaviors **(Table S2)**. This modelling allows the extraction of key parameters, such as the maximum fluorescence intensity (MFI), which corresponds to the total amyloid formation, and the half-time of aggregation (t_1/2_), used to assess the overall speed of the process. Our analysis revealed that hIAPP alone reached an <u>MFI of 14,100 a.u.</u>, while in the presence of QBP1 this value was drastically reduced to <u>1,560</u> <u>a.u.</u> **(Figure 1D)**. Moreover, t_1/2_ decreased significantly when incubated with QBP1, shifting from <u>4.4 h</u> for hIAPP alone to <u>0.49 h</u> in the presence of QBP1 (data not shown, extracted from **Table S2**). While a shorter t_1/2_ is generally associated with accelerated aggregation, in this case the consistently low fluorescence signal throughout the experiment indicates that QBP1 effectively inhibits fibril formation rather than promoting a faster aggregation process. Although the hIAPP: QBP1 curve may appear to exhibit a more rapid elongation phase, this interpretation is misleading due to the overall lower fluorescence intensity, which reflects a substantial reduction in amyloid formation. More importantly, the drastic reduction in fluorescence signal from the early stages suggests that QBP1 exerts its effect at the initial phases of aggregation, likely by interfering with monomeric hIAPP and its conformational transition into aggregation-prone β-sheet structures. This highlights the role of QBP1 in preventing the formation of early oligomeric species, which are crucial toxic precursors to mature amyloid fibrils.

To get further insight into the mechanism of action of QBP1, we tested two SC control variants of QBP1, SC-M8 and SC-M11 (Nagai et al., 2007, 2000). Surprisingly, although less effective than QBP1, both significantly inhibited amyloid formation **(Figure 1C)**. Non-linear modelling in OriginPro showed that hIAPP alone and in the presence of the SC variants followed a Boltzmann model **(Table S3)**, indicating similar aggregation kinetics. However, with SC treatment, hIAPP aggregation kinetics displayed an extended lag phase and a shortened elongation phase, suggesting reduced nucleation and fibril growth. Fluorescence intensity measurements revealed that hIAPP alone reached a maximum of 14,330 a.u., which dropped significantly with SC-M8 (8,849 a.u.) and SC-M11 (9,570 a.u.) **(Figure 1E)**. Additionally, t_1/2_ increased from 4.61 h (hIAPP alone) to 8.62 h (SC-M8) and 9.91 h (SC-M11), extracted from **Table S3**. These results indicate that although SC peptides delay amyloid formation and reduce total fibril accumulation, their inhibitory effect is less pronounced than that of QBP1. Therefore, the inhibitory activity of SC-M8 and SC-M11, despite lacking a defined conformation and hydrophobic core (Ramos-Martín et al., 2014), underscores the role of sequence-specific interactions in QBP1 action. Still, non-sequence-specific interactions, likely involving aromatic residues, also seem to contribute to reduce amyloid aggregation, emphasizing the importance of both composition as well as sequence and conformation in QBP1-mediated inhibition.

Finally, fluorescence microscopy (20×) was employed to visualize ThT-stained hIAPP samples after 24 and 48 h of incubation, with or without inhibitory peptides **(Figure 1F, Figure S2)**. ThT-positive aggregates were evident under all conditions, indicating an ongoing amyloid formation. However, in the hIAPP-alone sample, the aggregate density increased over time, with large, densely packed mesh-like structures already prominent at 24 h and further accentuated by 48 h. These assemblies exhibited irregular, branched morphologies consistent with amyloid networks (Pytowski et al., 2020; Shi et al., 2019). Due to the resolution limitations of fluorescence microscopy, their ultrastructural identity was further examined using TEM, which confirmed their amyloid fibrillar nature, as described below. In contrast, co-incubation with QBP1 (1:5 molar ratio) markedly reduced aggregate density and altered their morphology, suggesting effective inhibition and/or modulation of fibril assembly. Remarkably, samples treated with SC-M8 and SC-M11 displayed intermediate fluorescence intensities and aggregate densities—higher than in QBP1-treated samples but lower than in the hIAPP-alone condition—suggesting partial inhibitory activity. These observations are consistent with the kinetic profiles **(Figure 1C)**, reinforcing that while SC peptides retain some capacity to modulate aggregation, they are less effective than QBP1. Additional images acquired at 10× magnification, including control conditions, are provided in the supplementary material **(Figure S2)** for reference.

To get higher-resolution insights into the aggregate morphology, samples were also analyzed at 40× magnification using Ibidi μ-Slides **(Figure 1G)**. As at lower magnification, hIAPP-alone samples showed dense, amorphous ThT-positive aggregates with the highest abundance. Co-incubation with QBP1 altered the distribution—smaller, fragmented, and more diffuse aggregates—mirroring the reduced fluorescence and kinetic delay seen in ThT assays and 20× imaging. Small spherical assemblies in both conditions (*top insets, first and second columns*) may represent immature or off-pathway intermediates. Distinctively, SC-M8 and SC-M11 treatments produced more defined spherical or ring-like assemblies with sharply demarcated edges (**Figure 1G***, top insets, third and fourth columns*). Some of these aggregates displayed a dark core surrounded by a fluorescent ring, suggesting a more organized molecular arrangement and a possible restriction in fibril elongation. These structured assemblies were more dispersed compared to the amorphous clusters in hIAPP-alone samples, suggesting that SC peptides redirect hIAPP aggregation toward alternative, possibly non-fibrillar pathways. Remarkably, in the IAPP: SC-M11 condition, liquid droplet-like condensates (LLPS) also appeared at the interface between the soluble fraction and the drying edge of the sample (**Figure 1G***, bottom inset, fourth column*), suggesting phase separation (Pytowski et al., 2021, 2020) as part of a distinct aggregation mechanism. Additional confocal microscopy and 3D reconstructions **(Figure S3)** confirmed the persistence and morphological divergence of these aggregates over a 7-day period.

### QBP1 inhibits hIAPP aggregation by targeting early oligomeric species

To assess QBP1’s ability to inhibit the formation of early amyloidogenic intermediates, we performed immunodot blot analysis using two conformation-specific antibodies: A11, which recognizes prefibrillar oligomers (Kayed et al., 2003), and OC, which detects fibrillar oligomers and mature fibrils (Kayed et al., 2007). As shown in **Figure 2A**, hIAPP alone exhibited a progressive increase in A11 and OC immunoreactivities over time, reflecting a clear transition from oligomeric to fibrillar species. A11 immunoreactivity first appeared at 16 h, indicating the formation of early prefibrillar oligomers, followed by OC positivity at 48 h, marking the presence of mature amyloid fibrils. In contrast, co-incubation with QBP1 (1:5 molar ratio, hIAPP: QBP1) drastically reduced A11 and OC signals at all-time points, demonstrating strong inhibition of amyloid polymerization. QBP1 itself did not form detectable aggregates **(Figure S4)**, and the negative control (BSA) lacked immunoreactivity, confirming the specificity of A11 and OC binding to amyloid structures. The significant reduction in A11 signal at early stages strongly suggests that QBP1 exerts its inhibitory effect by preventing the assembly of early oligomeric precursors, which are essential for fibril maturation.

**Figure 2.**
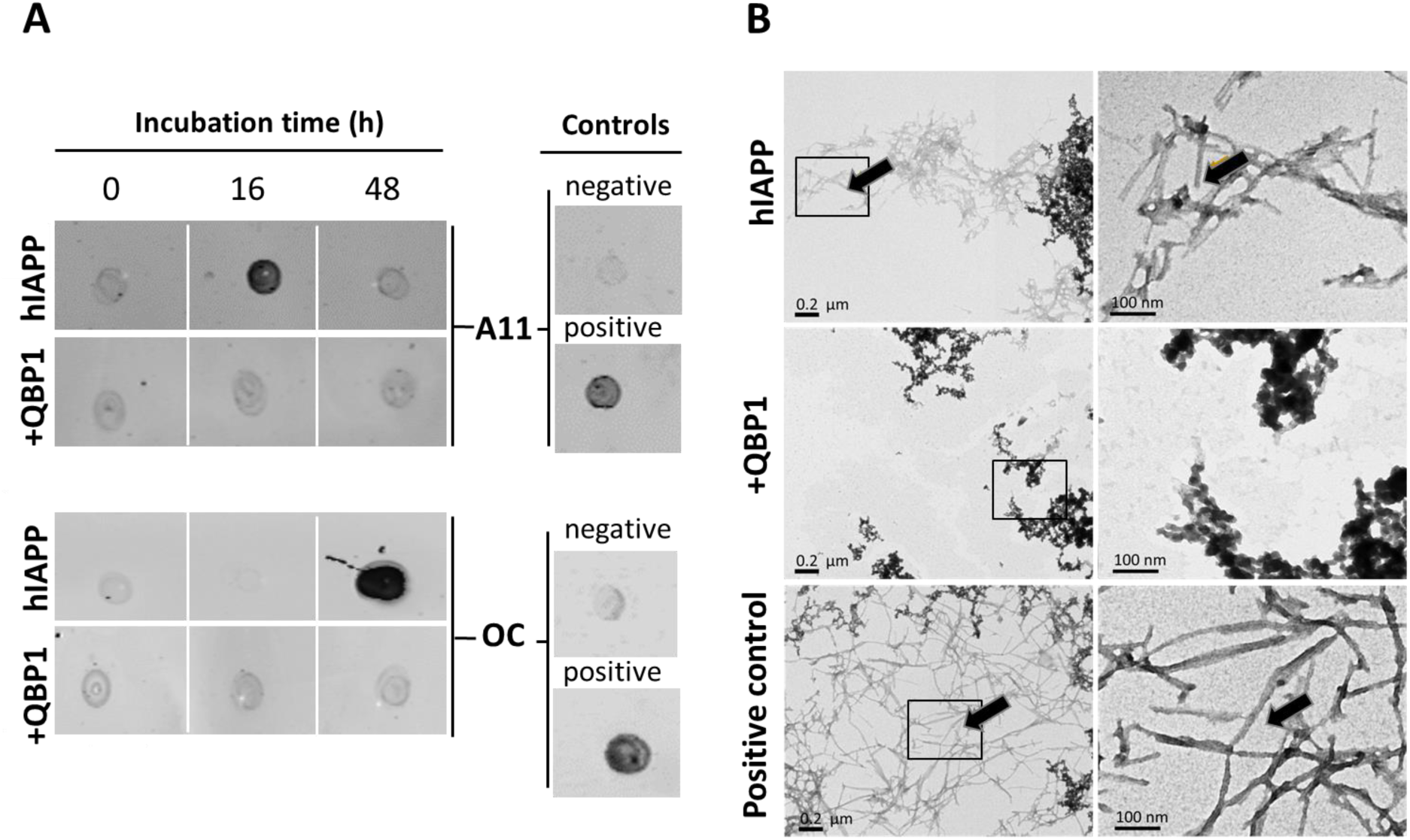
QBP1 disrupts hIAPP amyloid aggregation by preventing the formation of early intermediates and fibril maturation. **(A)** Immunodot-blot analysis of <u>hIAPP</u> (70 μM) alone or co-incubated with QBP1 (1:5 molar ratio) at 0, 16, and 48 h, using A11 (oligomer-specific) or OC (fibril-specific) antibodies. hIAPP alone showed increasing A11 and OC signals over time, while QBP1 co-incubation significantly reduced both, inhibiting oligomer and fibril formation. Controls: BSA (non-amyloidogenic, negative) and Aβ42 (amyloidogenic, positive). **(B)** Representative TEM images of hIAPP alone, hIAPP: QBP1 (1:5), and α-Syn fibrils (positive control) incubated for 72 h. hIAPP alone formed dense fibrillar networks (black arrows), whereas QBP1-treated samples showed fewer fibrils and only small, dispersed aggregates, suggesting non-specific protein clustering rather than amyloid polymerization. The insets within the 0.2 μm-scale images highlight regions selected for higher-magnification TEM at 100 nm resolution, providing a detailed view of the aggregate structures. Scale bars: 0.2 μm (left) and 100 nm (right). Representative images from three independent experiments.

To further evaluate the effects of prolonged incubation, samples were analyzed after 7 days in the presence of QBP1, SC-M8, and SC-M11 **(Figure S4)**. In all cases, A11 and OC signals were markedly reduced compared to hIAPP alone, indicating that both QBP1 and its SC variants effectively hindered amyloid progression. These findings reinforce the importance of both composition as well as sequence and conformation in QBP1-mediated inhibition. Interestingly, despite being ThT-positive, SC-induced aggregates **(Figure S3)** did not show A11 or OC immunoreactivity, suggesting that they lack the canonical prefibrillar or fibrillar conformations recognized by these antibodies. This may indicate that SC variants drive hIAPP aggregation into structurally distinct, non-classical assemblies, or that these aggregates are highly compact or sterically hindered, which would prevent antibody binding but not ThT intercalation.

Finally, to directly assess the structural effects of QBP1 on hIAPP aggregation, we employed TEM at 25,000× and 100,000× magnifications **(Figure 2B)**. In the hIAPP-alone condition, dense clusters of amyloid fibrils were observed, resembling those in the α-Synuclein positive control. In contrast, co-incubation with QBP1 significantly reduced fibril formation, resulting in only small, dispersed aggregates, the interpretation of which appears to be more consistent with non-specific protein clustering than with amyloid polymerization. This structural disruption suggests that QBP1 interferes with the ordered assembly of amyloid fibrils, likely by stabilizing early intermediates or off-pathway aggregates, thus preventing their conversion into fibrils.

### QBP1 protects INS-1E pancreatic β-cells from hIAPP-induced cytotoxicity

To assess the protective effect of QBP1 under conditions that mimic β-cell stress associated with hIAPP overexpression, we employed rat INS-1E pancreatic β-cells stably expressing human IAPP (INS-1E-hIAPP), along with non-transduced INS-1E cells as controls (INS-1E-Ct). Immunostaining confirmed hIAPP overexpression and efficient intracellular delivery of *Antp*-QBP1 **(Figure 3A)**. Indeed, hIAPP localized predominantly in the cytoplasm, with higher intensity in INS-1E-hIAPP cells compared to controls, reinforcing the validity of this model. To evaluate *Antp*-QBP1 internalization, cells were incubated with 10 μM peptide for 48 h, followed by extensive washing to remove non-internalized peptide. Subsequent immunofluorescence analysis revealed robust cytoplasmic uptake in nearly all β-cells, with minimal nuclear presence, indicating both stability and efficient intracellular delivery. Additionally, *Antp*-QBP1 showed extensive colocalization with hIAPP in INS-1E-hIAPP cells, consistent with their shared cytoplasmic distribution. While this observation does not confirm a direct interaction, it supports the possibility of spatial proximity that may facilitate QBP1’s inhibitory effect on hIAPP aggregation contributing to its protective role in β-cell function.

**Figure 3.**
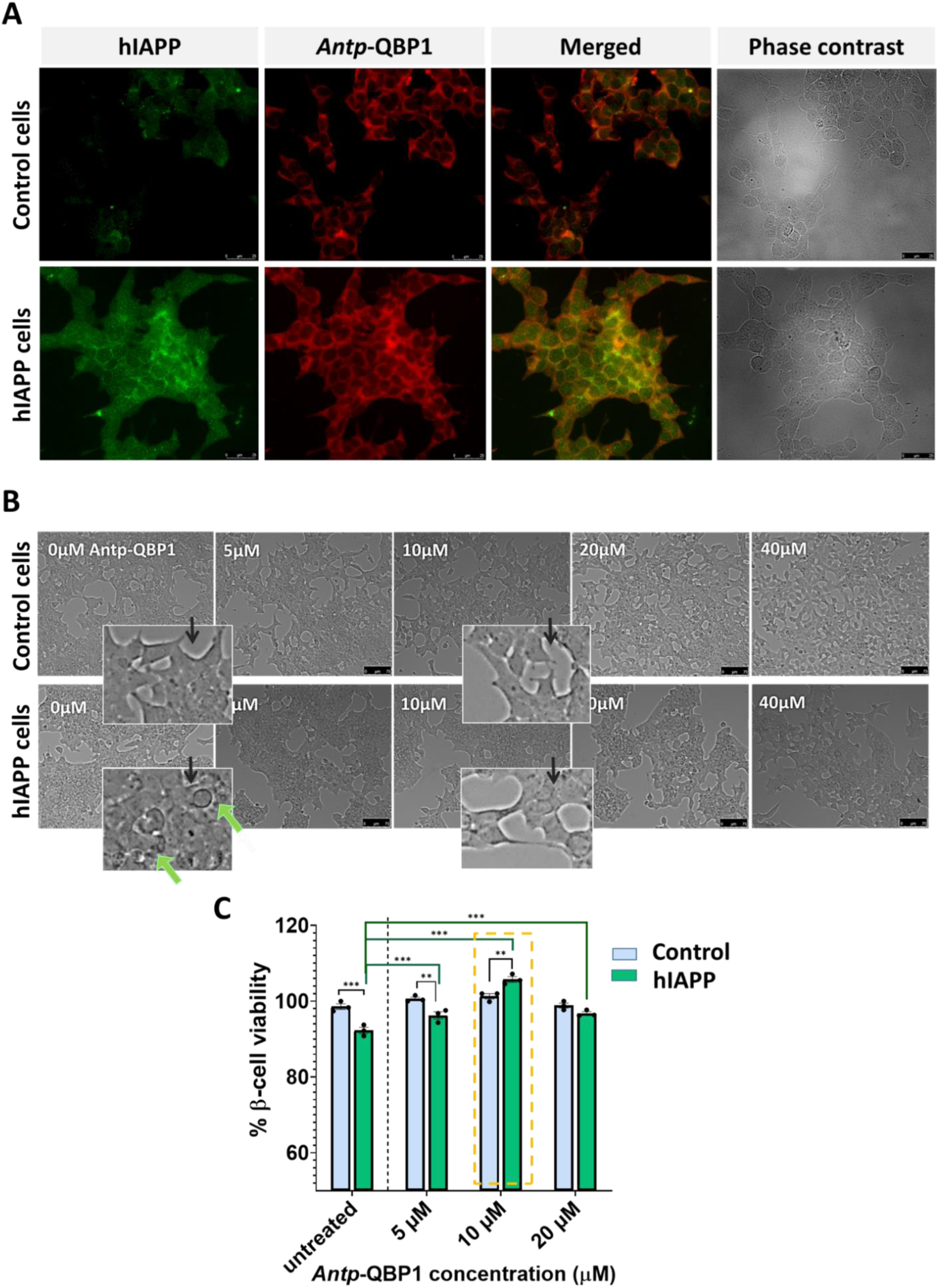
*Antp*-QBP1 facilitates the intracellular delivery of QBP1 and protects INS-1E β-cells from hIAPP-induced toxicity. **(A)** Immunofluorescence analysis of hIAPP (first column, green) and *Antp*-QBP1 (second column, red) in INS-1E-hIAPP and control cells after 48 h of exposure to 10 µM *Antp*-QBP1. Merged images (third column) show intracellular colocalization, suggesting potential spatial proximity between hIAPP and QBP1. Images were acquired using a Leica DMI 6000 Fluorescence Microscope (40× objective). **(B)** Phase-contrast microscopy images of INS-1E-hIAPP and control cells, untreated or treated with *Antp*-QBP1 (5–40 μM) for 48 h. In the absence of peptide (0 μM *Antp*-QBP1), INS-1E-hIAPP cells displayed signs of cytotoxic stress, including increased cytoplasmic density, rounding, loss of substrate adhesion, and detachment (highlighted by green arrows). Treatment with *Antp*-QBP1 improved cell morphology and restored adhesion. Insets show selected areas at higher magnification. Images were captured using a Leitz DM IRB microscope (10×/0.40 objective, K5 camera, Leica, Germany). Scale bars: 75 μm. Representative images are shown. **(C)** Quantification of β-cell viability in INS-1E-hIAPP and control cells after 48 h of *Antp*-QBP1 treatment (5–20 μM). hIAPP expression significantly reduced cell viability compared to control cells (p < 0.001). *Antp*-QBP1 treatment significantly improved viability at all concentrations tested (p < 0.001), when compared to untreated INS-1E-hIAPP cells. Notably, 10 μM *Antp*-QBP1 increased viability by ∼10% (p = 0.002), and 20 μM restored viability to control levels (not significant vs. controls). Data are presented as mean ± SEM from three replicates (CellTiter-Glo® assay). Statistical significance: *p < 0.05, **p = 0.002, ***p = 0.001; comparisons as indicated in the graph.

We next explored whether *Antp*-QBP1 could also counteract the detrimental effects of hIAPP overexpression on β-cell morphology and viability. Phase-contrast microscopy was employed to assess morphological changes after 48 h of *Antp*-QBP1 exposure **(Figure 3B)**. In the absence of treatment, marked morphological differences were evident between control and hIAPP-overexpressing cells. Specifically, INS-1E-hIAPP cells displayed classic signs of β-cell stress and dysfunction, including increased cytoplasmic density, rounding, loss of substrate adhesion, and detachment—reflecting a compromised state driven by hIAPP overexpression. Remarkably, treatment with increasing *Antp*-QBP1 concentrations (5–40 μM) led to clear morphological improvements. Cells treated with 10 or 20 μM showed substantial recovery, with reduced rounding and detachment, and improved adhesion—closely resembling untreated control cells.

To determine whether these changes correlated with enhanced cell survival, β-cell viability was assessed under the same conditions. Both INS-1E-Ct and INS-1E-hIAPP cells were treated with 5–20 μM *Antp*-QBP1 for 48 h, and viability was quantified **(Figure 3C)**. Consistent with previous reports of hIAPP-induced cytotoxicity (Haataja et al., 2008), hIAPP-expressing cells showed a significant reduction in viability compared to controls (p < 0.001), confirming the deleterious effect of hIAPP overexpression. Importantly, *Antp*-QBP1 treatment significantly improved INS-1E-hIAPP cell viability at all tested concentrations (p < 0.001), highlighting its protective effect. Notably, at 10 µM, *Antp*-QBP1 led to a ∼10% increase in cell viability (p < 0.002) compared to untreated hIAPP-expressing cells, nearly restoring viability levels to those of control cells treated with the same dose of QBP1 (*highlighted in Figure 3C, yellow dashed box*). At 20 µM, *Antp*-QBP1 fully restored viability to levels comparable to control cells, further reinforcing its protective role. These results strongly suggest that QBP1 mitigates the cytotoxic effects of hIAPP overexpression, enhancing β-cell survival.

While these findings align with ThT aggregation data and support a protective role for QBP1 against hIAPP-induced cytotoxicity, they do not conclusively demonstrate that amyloid inhibition is the primary mechanism involved. Alternative mechanisms—such as disruption of hIAPP interactions or modulation of cellular stress responses—may also contribute. To clarify the role of hIAPP amyloidogenesis in cytotoxicity and further elucidate the protective mechanism of *Antp*-QBP1, we employed a more controlled experimental setup in which INS-1E β-cells were exposed to <u>exogenous hIAPP oligomers</u>. This approach allowed precise control over hIAPP concentrations and enabled a systematic assessment of both dose-dependent effects and the impact of QBP1 treatment on β-cell viability, morphology, and gene expression **(Figure 4)**.

**Figure 4.**
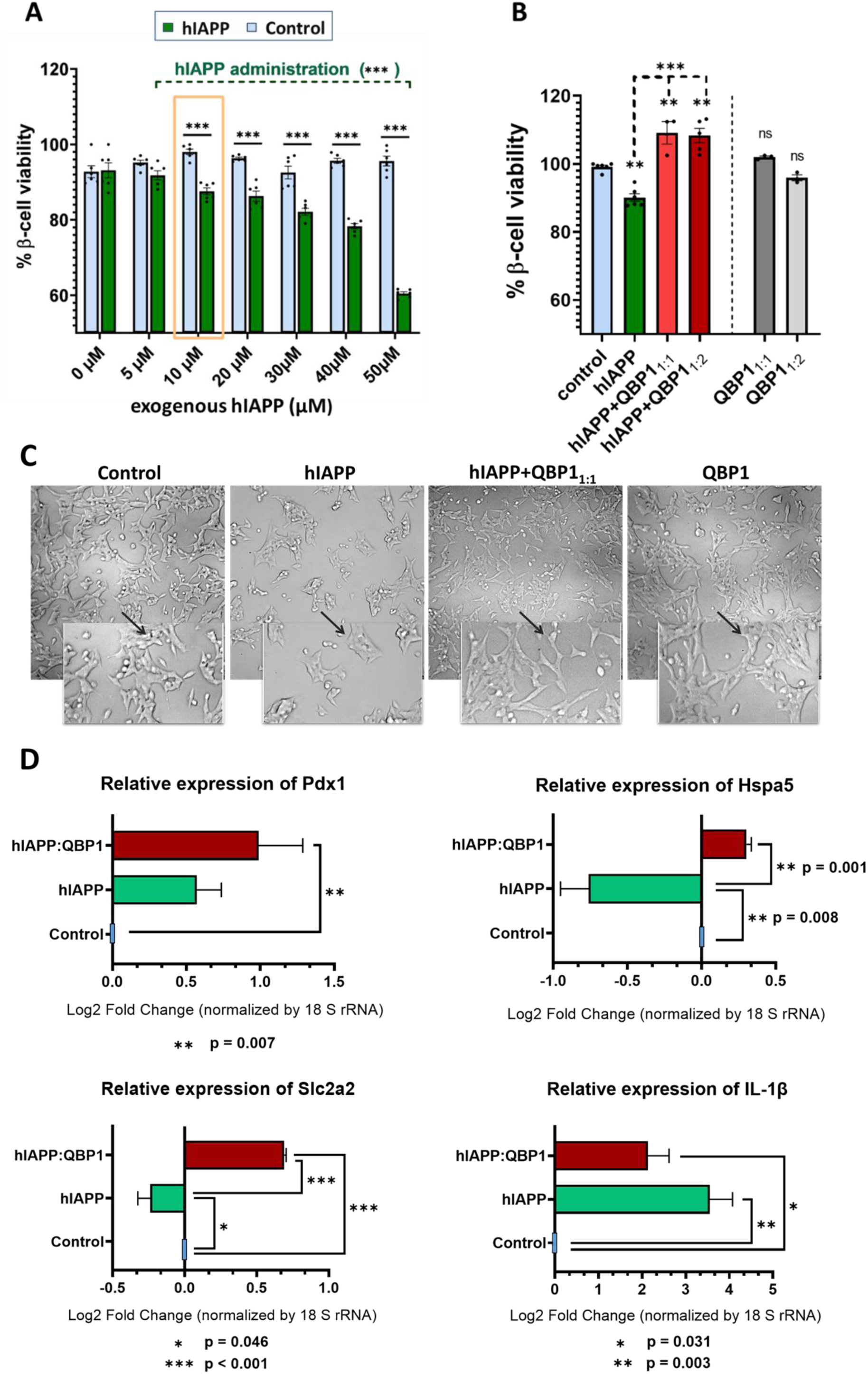
QBP1 protects INS-1E β-cells from hIAPP oligomer-induced toxicity by preserving viability and cellular integrity. (A) Cell viability assay: INS-1E β-cells were exposed to increasing concentrations of exogenous hIAPP oligomers (5–50 μM) for 48 h, showing a dose-dependent viability reduction (p < 0.001), while 5 μM showed no significant effect. The yellow dashed box highlights the 10 μM hIAPP as the minimum concentration significantly reducing viability (p < 0.001). **(B) Protective effect of QBP1:** Co-incubation of 10 μM hIAPP with *Antp*-QBP1 (1:1 and 1:2 ratios) significantly improved viability (p < 0.001 vs. hIAPP alone), confirming its cytoprotective effect. While QBP1 alone (1:1 or 1:2) had no significant effect (ns). Data are plotted as mean ± SEM (CellTiter-Glo® assay). Statistical significance: *p < 0.05, **p = 0.01, ***p = 0.001; ns = not significant; comparisons as indicated in the graph. **(C) Cell morphology analysis:** Phase-contrast microscopy images after 48 h of treatment. hIAPP exposure led to a marked reduction in cell confluence and subtle morphological alterations (middle panel), while *Antp*-QBP1 co-treatment (right panel) preserved morphology and adhesion, resembling the control condition (left panel). Insets show detailed structures. Images were captured using a Leitz DM IRB microscope. Scale bars: 75 μm. Representative images are shown. **(D) Gene expression analysis:** Relative mRNA expression of key β-cell identity and stress-related genes after 48 h treatment, normalized to 18S rRNA. Exogenous hIAPP (10 μM) significantly downregulated *Slc2a2* and *Hspa5* while upregulated *IL-1β* in INS-1E β-cells, with no significant effect on *Pdx1*. QBP1 co-treatment restored *Slc2a2* and *Hspa5* expression, reduced *IL-1β* levels, and upregulated *Pdx1*, suggesting enhanced β-cell function, endoplasmic reticulum homeostasis, and reduced inflammation. Data are plotted as mean ± SD (n = 3); statistical significance was determined using Bonferroni’s post-hoc test for multiple comparisons (p < 0.05 considered significant, exact p values shown in graph).

Since hIAPP oligomers are widely recognized as the major toxic species, INS-1E β-cells were cultured overnight in high-glucose medium (16.7 mM), and then exposed to increasing concentrations (5–50 μM) of extracellular hIAPP oligomers for an additional 48 h, either alone or in combination with *Antp*-QBP1. To ensure the presence of prefibrillar oligomers, A11-positive hIAPP samples—preincubated to promote oligomerization—were sonicated before being added to the culture medium. As shown in **Figure 4A**, hIAPP oligomers induced a dose-dependent decline in β-cell viability (p < 0.001), with significant effects observed from 10 μM onwards, confirming their cytotoxic potential. In contrast, 5 μM hIAPP did not significantly affect viability, suggesting that this concentration is subthreshold under our experimental conditions.

Based on these results, 10 μM—being the lowest concentration to significantly reduce viability (p < 0.001)—was selected to evaluate whether *Antp*-QBP1 could prevent β-cell damage during the early stages of oligomer formation. To target the early stages of amyloidogenesis, *Antp*-QBP1 was preincubated with hIAPP before treatment and then simultaneously introduced into the culture medium. As expected, hIAPP alone significantly reduced cell viability compared to the PBS control (p < 0.01). However, the addition of *Antp*-QBP1 at either an equimolar ratio (1:1) or a two-fold molar excess (1:2) not only prevented viability decline but also enhanced survival, resulting in an approximately 19% increase compared to the hIAPP-alone group (p < 0.001). These findings suggest that *Antp*-QBP1 prevents or even reverses hIAPP aggregation, thereby mitigating β-cell toxicity. Notably, increasing *Antp*-QBP1 from 10 to 20 μM did not further enhance the effect, suggesting a potential saturation point in its activity. This plateau effect may be due to saturable binding kinetics or solubility limitations affecting molecular interactions. Further investigations are warranted to elucidate these dose-dependent effects and the underlying molecular mechanisms. Moreover, QBP1 alone (1:1 or 1:2) did not significantly affect β-cell viability compared to the PBS control (ns, not significant), suggesting that QBP1 does not promote β-cell proliferation under these conditions **(Figure 4B)**.

We next examined whether *Antp*-QBP1 could also mitigate the effects of exogenous hIAPP on β-cell morphology. Phase-contrast microscopy was used to assess morphological changes after 48 h of treatment **(Figure 4C)**, following the procedure described in **Figure 3B**. In contrast to the acute cytotoxic features seen in earlier models **(Figure 3B)**, hIAPP exposure here primarily led to a marked reduction in cell confluence and subtle morphological alterations, with cells appearing more rounded or amorphous and lacking the elongated, spindle-like shape characteristic of healthy INS-1E β-cells. These observations suggest that hIAPP oligomers not only induce toxicity but also impair β-cell proliferation or survival over time. Notably, co-treatment with *Antp*-QBP1 preserved both confluence and morphology, with cells displaying a more organized, adherent, and elongated appearance, closely resembling the control untreated condition. Moreover, QBP1 alone did not induce detectable changes in cell morphology, reinforcing its role as a protective factor rather than a stressor.

### QBP1 modulates transcriptional responses to hIAPP aggregation, restoring β-cell function and resilience

To identify key molecular pathways affected by hIAPP aggregation and QBP1 treatment, we analysed differential gene expression between treatments using RQ-PCR. Four essential **β-cell regulatory genes associated with diabetes** were selected: ***Pdx1*** *(Pancreatic and duodenal homeobox 1)*, essential for β-cell identity and insulin gene transcription; ***Slc2a2*** *(Solute carrier family 2, facilitated glucose transporter member 2, GLUT2)*, a key glucose transporter involved in insulin secretion; ***Hspa5*** *(Heat shock protein family A (Hsp70) member 5, BiP/GRP78)*, a molecular chaperone regulating endoplasmic reticulum (ER) homeostasis and proteostasis; and ***IL-1β*** *(Interleukin-1 beta)*, a pro-inflammatory cytokine implicated in β-cell dysfunction and amyloid-induced toxicity (Back and Kaufman, 2012; Casas et al., 2007; Rodríguez-Comas et al., 2020). Gene expression was normalized to 18S rRNA **(Table S1)**, and changes in hIAPP-treated cells compared to controls, with and without QBP1 co-treatment were assessed **(Figure 4D)**.

hIAPP exposure for 48 h resulted in distinct transcriptional alterations. Thus, *Slc2a2* and *Hspa5* expression were significantly downregulated (*p* = 0.046 and *p* = 0.008, respectively), suggesting impaired glucose uptake and dysregulated ER stress responses, both of which are critical for β-cell survival. Additionally, *IL-1β* was markedly upregulated (*p* = 0.003), reinforcing its role in hIAPP-induced inflammatory stress. In contrast, *Pdx1* expression remained unchanged in the hIAPP condition, indicating that hIAPP alone does not strongly impact β-cell transcriptional regulation. However, QBP1 co-treatment reversed these detrimental effects. Thus, *Slc2a2* and *Hspa5* were significantly upregulated (*p* < 0.001), restoring glucose transporter expression and ER homeostasis, while *IL-1β* levels were reduced (*p* = 0.031), suggesting an attenuation of hIAPP-induced inflammation. Moreover, *Pdx1* expression was significantly increased with QBP1 (*p* = 0.007), reinforcing its role in maintaining β-cell identity and function. These findings suggest that hIAPP aggregation disrupts key β-cell pathways, impairing metabolic function and ER stress adaptation as well as increasing inflammatory stress, whereas QBP1 exerts a protective effect by enhancing β-cell resilience through transcriptional regulation of glucose transport, ER homeostasis, and inflammatory pathways. Further studies are required to determine whether these molecular adaptations translate into long-term improvements in β-cell survival and insulin secretion under amyloidogenic stress.

### Molecular insights into the QBP1-hIAPP interaction through MD simulations

To explore the nature of the binding interactions between **amylin** (**PDB ID: 2L86** (Nanga et al., 2011); depicted in **Fig. S1**) and the peptides QBP1 (Ramos-Martín et al., 2014), SC-M11 and SC-M8 (Popiel et al., 2007; Ramos-Martín et al., 2014), we performed MD simulations. **Figure 5A** summarizes the main structural and physicochemical features associated to the hIAPP sequence as well as to the inhibitory peptides, based on previous reports. hIAPP has a highly hydrophobic core, particularly in its central region, which promotes self-aggregation and fibril formation (Bakou et al., 2017; Fortier et al., 2022). Additionally, its C-terminal region plays a crucial role in modulating aggregation, primarily due to the presence of aromatic and polar residues that influence intermolecular interactions. In particular, Tyrosine (Y37) can engage in π-π interactions and hydrogen bonding, potentially affecting oligomerization and amyloid assembly, although its role in fibril stabilization remains unclear (Caillon et al., 2016; Cao et al., 2020). QBP1 displays a well-defined hydrophobic core, composed primarily of tryptophan (W), phenylalanine (F), and proline (P) residues (Ramos-Martín et al., 2014), which may contribute to its interaction with hIAPP’s hydrophobic domains. The presence of glycine (G) enhances flexibility, while P may interfere with amyloid aggregation (Ghosh et al., 2020; Nayak et al., 2024), potentially improving its inhibitory activity. In contrast, SC-M11 and SC-M8 have rearranged hydrophobic and aromatic residues, possibly altering their ability to establish π-π interactions and hydrogen bonds with hIAPP, affecting their inhibitory efficiency (Ramos-Martín et al., 2014).

**Figure 5.**
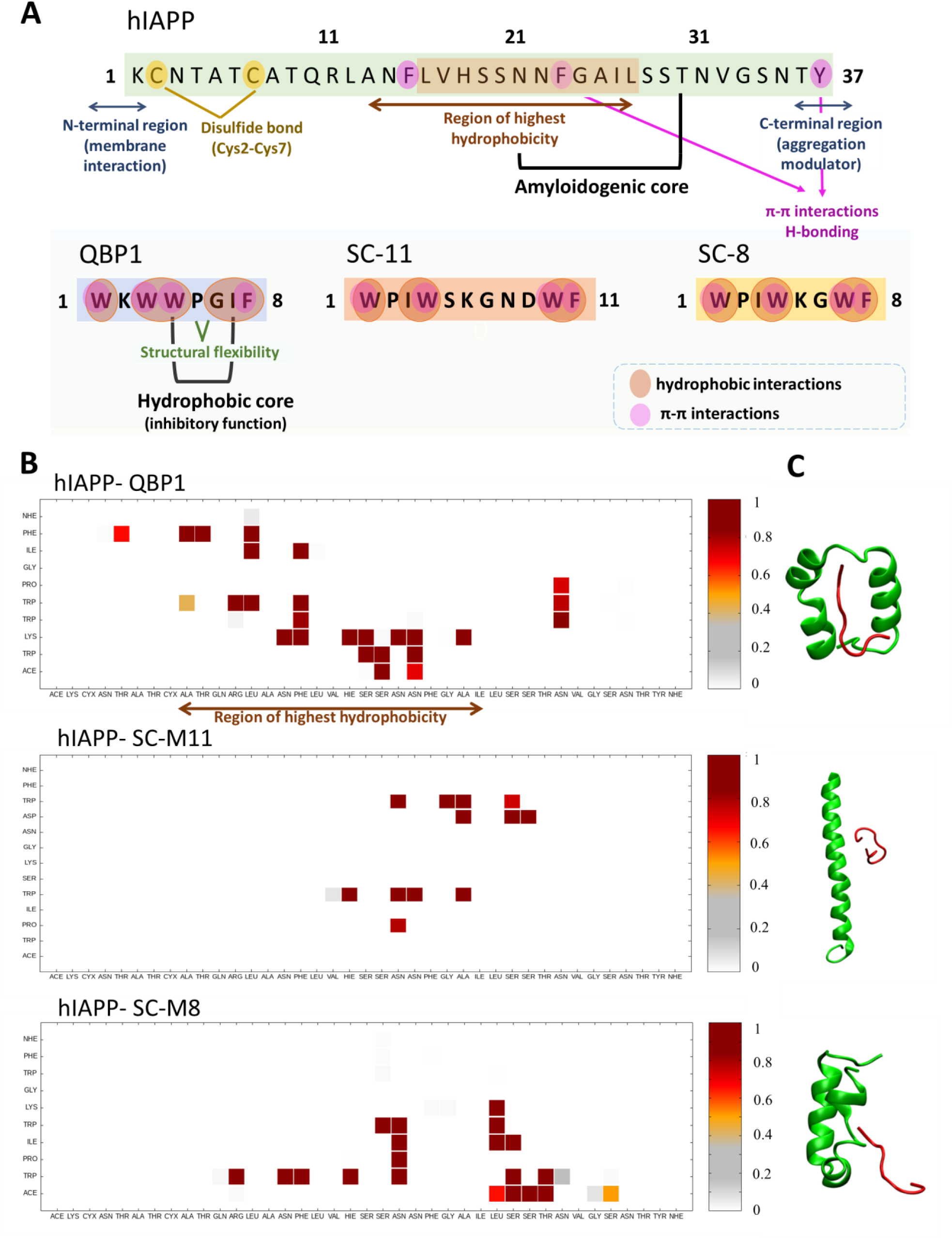
Stable binding modes and contact interactions between amylin and QBP1, SC-M11, and SC-M8. **(A)** Schematic representation of the primary structural and physicochemical features of hIAPP and the inhibitory peptides. The hydrophobic core of hIAPP, key aromatic residues, and regions involved in intermolecular interactions are highlighted, illustrating their potential role in self-aggregation and inhibition. The inhibitory peptides are depicted with their respective hydrophobic and flexible regions, which may influence their binding to hIAPP. **(B)** Contact maps illustrating the most frequently observed interactions in the amylin-QBP1 (top panel), amylin-SC-M11 (middle panel), and amylin-SC-M8 (bottom panel) complexes across the MD simulations. The color scale represents the probability of contact formation, with red indicating higher probabilities and grey indicating lower probabilities. CYX denotes CYS residues forming a disulfide bridge, which may contribute to structural stability. **(C)** Structural representations of the amylin-QBP1 (top panel), amylin-SC-M11 (middle panel), and amylin-SC-M8 (bottom panel) complexes, highlighting the most common contact regions. Amylin is shown in green, while the inhibitors are depicted in red. The structural differences among the complexes highlight variations in binding interactions and stability, with QBP1 forming a more compact and stable interaction network compared to its scrambled variants.

To assess the possible interactions involved, we calculated the contact map averaged over all frames of our simulations, obtaining a pattern that represents the most probable binding mode **(Figure 5B)**. From the most probable and stable structures of each binding mode, retrieved as described in Material and Methods section, complex stability and affinity through binding energy calculations are listed **(Table 1)**. By comparing QBP1 with its SC variants, we have identified possible key molecular determinants governing amylin recognition and inhibition, which should provide insights into the structural and energetic factors underlying these interactions.

**Table 1.** Binding energy components of QBP1, SC-M11, and SC-M8 complexes. The table shows the energy contributions to the binding free energy (*ΔGbind*) for the amylin-QBP1, amylin-SC-M11, and amylin-SC-M8 complexes, calculated from molecular dynamics simulations. The components include: van der Waals interactions (*ΔE_vdW_*), reflecting non-covalent attractive forces; Electrostatic interactions (*ΔE_ele_*), capturing charge-charge interactions; Polar solvation energy (*ΔG_PB_*) and non-polar solvation energy (*ΔG_NP_*), representing solvent effects on binding. Additionally, entropic contributions (*−TΔS*) and total binding free energy (*ΔG_bind_*), and *enthalpy* values are provided. In this context, Enthalpy represents the binding energy excluding entropic effects. All values are given in kcal/mol, with more negative values indicating stronger contributions to binding stability.

| | $\Delta E_{vdW}$ | $\Delta E_{ele}$ | $\Delta G_{PB}$ | $\Delta G_{NP}$ | $-T\Delta S$ | $\Delta G_{bind}$ | Enthalpy |
| --- | --- | --- | --- | --- | --- | --- | --- |
| *QBP1* | -55.487 | 2.718 | 23.464 | -6.205 | 29.840 | -5.704 | -35.544 |
| *SC-M11* | -31.271 | -0.294 | 12.673 | -3.068 | 18.180 | -3.793 | -21.982 |
| *SC-M8* | -39.640 | -1.515 | 15.457 | -3.997 | 26.175 | -3.520 | -29.713 |

The most stable binding modes, identified from MD simulation trajectories of amylin-QBP1, amylin-SC-M11, and amylin-SC-M8 complexes, were characterized by identifying the frequency of residue-residue contacts in the simulations **(Figure 5B)**, which would translate to the probability of these pairs forming an actual contact upon interaction. The amylin-QBP1 complex exhibited high-probability interactions with hydrophobic and aromatic-rich regions (*e.g.*, W, F) **(Figure 5B, Figure S5)**. These contacts (shown in as a range of red tones in the contact maps) indicate a stable binding pattern that helps maintain amylin’s structure and prevent aggregation. Additionally, the disulfide bridge between Cys2 and Cys7 in hIAPP may contribute to overall structural stability, indirectly supporting the inhibitory effect of QBP1. Importantly, QBP1 exhibits its strongest interactions with key amyloidogenic regions of amylin—residues 8–20, 20–29, and 30–37—which are critical for β-sheet formation. By targeting these regions, QBP1 likely interferes with fibrillogenic pathways. In contrast, SC-M11 and SC-M8 showed weaker and lower-probability interactions, suggesting suboptimal binding motifs. Although SC-M8 displayed moderate binding potential, both SC peptides were notably less effective than QBP1 in stabilizing amylin and inhibiting hIAPP aggregation. These findings are further supported by the averaged contact maps **(Figure S6)**, which complement this analysis by showing the most probable contacts formed across all simulations, reinforcing the consistency of the identified binding regions.

To quantify the binding strength, binding free energy calculations were performed using MM/PBSA method **(Table 1)**. QBP1 exhibited the most favorable binding free energy (<u>ΔG_bind_ = −5.70 kcal/mol</u>), indicating a higher affinity for amylin compared to <u>SC-M11 (−3.79 kcal/mol</u>) and <u>SC-M8 (−3.52 kcal/mol</u>) (Copeland, 2013; Pantsar and Poso, 2018). These findings are consistent with the lower interaction stability of SC-M11 and SC-M8 observed in contact maps **(Figure 5B)**. Furthermore, QBP1 adopts a compact and well-aligned conformation that optimizes its interaction with amylin, whereas the SC peptides show reduced structural complementarity and stability **(Figure 5C)**. This suggests that the superior stability of QBP1 arises not only from its favorable binding energy, but also from its ability to effectively engage key amyloidogenic regions of the target peptide.

To gain deeper insights into the nature of these interactions, the total binding energy was further decomposed into its individual energy components **(Table 1, Figure S7)**. The stabilizing effect of hydrophobic interactions in the QBP1–amylin complex was especially pronounced, with van der Waals forces (ΔE_vdW_ = 055.49 kcal/mol) emerging as the dominant contributor. This underlines the critical role of W and F residues in mediating complex formation. In contrast, electrostatic contributions (ΔE_ele_) were negligible in SC-M11 (−0.294 kcal/mol) and SC-M8 (−1.515 kcal/mol), indicating that Coulombic interactions did not significantly favor binding in these peptides. Solvation effects also played a critical role in overall binding stability. Although polar solvation energy (ΔG_PB_) was unfavourable across all complexes—most notably for QBP1 (23.464 kcal/mol)— this desolvation cost was counterbalanced by its strong hydrophobic stabilization (ΔG_NP_ = −6.205 kcal/mol). SC-M11 and SC-M8 exhibited lower desolvation costs (12.673 kcal/mol and 15.457 kcal/mol, respectively) but also weaker hydrophobic stabilization (ΔG_NP_ = −3.068 kcal/mol and −3.997 kcal/mol, respectively), leading to an overall reduction in binding stability compared to QBP1. This energetic pattern suggests that peptide– amylin interactions are predominantly driven by hydrophobic forces while inefficiently shielding polar groups from solvent exposure—a behavior characteristic of partially misfolded or intermediate aggregation states. Such interactions also align with a tendency toward LLPS, both possibilities consistent with our findings.

In addition, specific aromatic interactions also appeared to contribute to complex stability. Notably, π-H bonds—stabilizing interactions between aromatic rings and hydrogen atoms, previously proposed in the QBP1–TDP-43 interaction (Mompeán et al., 2019)—were more prevalent in QBP1 than in SC-M11 and SC-M8 **(Figure S8)**. These aromatic interactions, together with the compact and well-aligned binding conformation of QBP1, likely contribute to its superior stability and inhibitory efficiency against amylin aggregation. Conversely, the lower frequency of π-H bonds observed in SC-M11 and SC-M8 may further explain their weaker binding affinities and reduced anti-amyloidogenic potential.

Taken together, the integration of binding energy decomposition, molecular interaction analysis, and structural observations highlights the superior stability and inhibitory efficiency of QBP1 over its SC variants. Strong van der Waals contributions, favorable non-polar solvation energy, and a higher prevalence of π-H bonds collectively enhance its affinity for amylin, reinforcing its potential as a therapeutic inhibitor of amylin aggregation. In contrast, SC-M11 and SC-M8 exhibit weaker binding, which seems to be primarily due to reduced van der Waals stabilization, less effective hydrophobic interactions, and unfavourable solvation effects. These findings provide a detailed molecular basis for the effectiveness of QBP1, supporting its potential role in mitigating amylin aggregation and its associated cytotoxic effects and pave the way for future peptide improvement by rational design if needed.

## Discussion

Accumulating evidence suggests that aggregates of hIAPP contribute significantly to pancreatic β-cell dysfunction and the progression of T2D (Kiriyama and Nochi, 2018; Mukherjee et al., 2015; Rambaran and Serpell, 2008). Due to its intrinsically amyloidogenic properties, hIAPP is prone to misfolding and aggregation, particularly under conditions of increased secretory demand (Akter et al., 2016; Haataja et al., 2008). These toxic species disrupt cellular homeostasis, trigger oxidative stress, and promote inflammatory responses, ultimately culminating in β-cell apoptosis and impaired insulin secretion (Cooper et al., 1988; Suárez et al., 2023). Elucidating these pathogenic mechanisms is therefore essential for the development of targeted therapies that can prevent or reverse hIAPP aggregation and potentially delay disease progression (Hassan et al., 2022). Although numerous aggregation inhibitors, including short peptides, have been investigated (Wirth et al., 2023; Xu et al., 2022), only a few have reached clinical approval, highlighting the urgent need for drug candidates with enhanced pharmacological properties. Among approved therapies, Symlin has been shown to reduce amyloidogenicity, limit aggregation, and decrease surface adhesion (Alrefai et al., 2010; McQueen and Bonk, 2005), but its clinical utility remains limited due to high production costs, poor solubility, and low oral bioavailability (Alrefai et al., 2010). In contrast, newer amylinomimetics such as Davalintide show greater potential, offering improved efficacy and extended activity (Boyle et al., 2022), highlighting the potential of optimized short peptides. Nevertheless, Davalintide remains in preclinical evaluation, and no other peptide-based therapies have yet achieved regulatory approval, reinforcing the need to advance promising candidates beyond early development.

Here, we have demonstrated that the QBP1 inhibitor, in particular its minimal active core (Nagai et al., 2000; Ren et al., 2001; Tomita et al., 2009), effectively disrupts early hIAPP amyloidogenesis, thereby protecting INS-1E β-cells from hIAPP-induced toxicity while preserving their viability and function. By stabilizing hIAPP through strong van der Waals interactions and π-H bonds at key hydrophobic and aromatic residues, QBP1 appears to prevent misfolding and reduce the formation of toxic oligomers and fibrils, mitigating downstream cytotoxic effects. This dual mechanism, primarily attributable to its high tryptophan content and robust hydrophobic interactions (Mompeán et al., 2019; Ramos-Martín et al., 2014; Tomita et al., 2009), is consistent with previous studies showing its ability to inhibit amyloid aggregation by targeting early toxic conformational transitions in various pathological models (Hervás et al., 2016; Nagai et al., 2007, 2003; Popiel et al., 2009, 2007; Yang et al., 2018). Other inhibitors, such as resveratrol and insulin, similarly act at early aggregation stages: resveratrol binds histidine (H18) to prevent oligomer formation, while insulin interacts with H18 and Y37 under acidic conditions to stabilize hIAPP (Lolicato et al., 2015; Wei et al., 2011). These findings reinforce the therapeutic relevance of early intervention strategies in amyloid-related pathologies. Taken together, our findings highlight QBP1 as a promising lead compound for preventing islet amyloid deposition in T2D (Kanatsuka et al., 2018; Potter et al., 2009).

Using ThT fluorescence spectroscopy and microscopy, we have observed that administering QBP1 at a 5-fold molar excess effectively inhibits hIAPP amyloidogenesis *in vitro* by reducing the formation of amyloid-like fibrils right from the earliest stages (Abedini et al., 2016; Ren et al., 2017). This inhibition implies that QBP1 interferes with the critical nucleation and elongation phases of aggregation (Ren et al., 2018; Shi et al., 2019; Young et al., 2017), thereby targeting the process at the earliest stage, before harmful species are formed. To determine whether QBP1 could prevent the formation of early-phase intermediates, known to be among the most cytotoxic species in the amyloid pathway (Kanatsuka et al., 2018; Soty et al., 2011), we performed conformational antibody assays using anti-oligomer A11 (Kayed et al., 2003) and anti-fibril OC (Kayed et al., 2007), which revealed a significant reduction in both oligomeric and fibrillar species. This suggest that QBP1 disrupts the assembly of early oligomeric precursors essential for fibril maturation (Chiti and Dobson, 2006). TEM further corroborated these findings by showing fewer hIAPP fibrils and a predominance of smaller, more dispersed aggregates, indicative of non-specific protein clustering, in the presence of QBP1 (Niu et al., 2020). Still, it must be noted that QBP1 did not completely prevent fibril formation, a limitation also observed with other peptide inhibitors such as SNNFGA and GAILSS, which similarly fail to entirely block hIAPP fibril formation (Kapurniotu et al., 2002; Scrocchi et al., 2002; Shi et al., 2019). Overall, these results indicate that QBP1 primarily delays the transition of hIAPP from its monomeric state to toxic oligomeric nuclei, consistent with its action on other amyloidogenic proteins (Hervás et al., 2012; Mompeán et al., 2019; Popiel et al., 2013).

Beyond its *in chemico* anti-amyloidogenic activity, QBP1 protected INS-1E-hIAPP β-cells from hIAPP-induced cytotoxicity, enhancing their survival, adhesion, and morphology (Hohmeier et al., 2000; Soty et al., 2011). These effects were supported by efficient intracellular delivery of the *Antp*-QBP1 peptide (Popiel et al., 2009, 2007). When compared to other delivery methods, such as viral vector-mediated gene delivery and liposomes, this PTD system offers several advantages, including notably high *in vivo* efficiency, low toxicity, and controllable administration (Popiel et al., 2011, 2007). Importantly, no cytotoxicity was observed up to 100 µM (Popiel et al., 2007), reinforcing its established safety profile (Bauer et al., 2010; López-García et al., 2025; Nagai et al., 2003).

We also confirmed hIAPP-induced cytotoxicity in our cell model, in contrast to some previous reports using the same INS-1E-hIAPP cells, where no significant increase in cell death was detected by annexin V/propidium iodide staining and flow cytometry (Soty et al., 2011). These discrepancies may stem from methodological differences or may reflect that hIAPP overexpression induces primarily functional impairments—such as reduced hormone secretion due to altered K-ATP channel activity—rather than overt apoptosis. In this context, increased mitochondrial metabolism has been proposed as a compensatory response to secretory defects (Soty et al., 2011). However, our findings are in line with those of Haataja *et al*. (2008), who reported hIAPP-induced apoptosis in COS-1 cells, further validating the cytotoxic potential of hIAPP and the relevance of our model. While INS-1E-hIAPP cells replicate key aspects of T2D pathology, amyloid deposition in these models typically requires higher hIAPP concentrations than those observed in diabetic patients (Alrouji et al., 2023; Soty et al., 2011). Moreover, *in vivo* amyloid formation is influenced by complex microenvironmental factors that are not fully reproduced *in vitro*. It must be noted that unlike traditional rodent models, which do not naturally develop amyloid deposits (Matveyenko and Butler, 2006), transgenic mice with stable hIAPP overexpression—particularly under a high-fat diet—develop amyloid deposits and more accurately mimic T2D progression (De Pablo et al., 2021; Montane et al., 2017). Therefore, the next step in our research will be to evaluate the therapeutic potential of QBP1 in these *in vivo* models, which will allow us to assess its efficacy under more physiologically relevant conditions.

To further investigate the contribution of hIAPP amyloidogenesis to β-cell dysfunction and validate the protective effect of QBP1, we performed complementary experiments using exogenously added hIAPP aggregates (Casas et al., 2007). This strategy enabled precise control of peptide concentration, allowing systematic assessment of dose-dependent toxicity and the impact of QBP1 on β-cell viability, morphology, and gene expression (De Pablo et al., 2021; Raleigh et al., 2017). It is well established that early-stage hIAPP oligomers—predominant during the lag phase of amyloid formation—are the most cytotoxic species, whereas mature fibrils are relatively inert (Abedini et al., 2016; Kanatsuka et al., 2018; Wirth et al., 2023). Consistent with this, A11 immunoreactivity confirmed the presence of toxic oligomers, which correlated with a dose-dependent decline in INS-1E β-cell viability. Based on our results and previous reports (Kanatsuka et al., 2018), 10 μM hIAPP was identified as the lowest concentration with a measurable toxic effect, and used as the threshold to assess the cytoprotective potential of *Antp*-QBP1. In addition, to mimic the hyperglycemic conditions of T2D, cells were cultured in high-glucose medium (16.7 mM), known to increase hIAPP expression and promote amyloid formation (Back and Kaufman, 2012; Lin et al., 2019). Under these conditions, *Antp*-QBP1—at equimolar or 2-fold molar excess—significantly preserved β-cell viability and restored normal cell adhesion and morphology. Notably, increasing the peptide concentration beyond this range did not further enhance its effect, suggesting saturable binding or solubility limitations (Kirsch et al., 2019; Lu et al., 2019). Together, these findings support the pathological relevance of extracellular hIAPP-induced toxicity in β-cells and confirm the protective capacity of QBP1, in line with previous models linking hIAPP aggregates to ER stress and β-cell failure (Casas et al., 2007).

Considering the complexity of hIAPP toxicity, which involves reactive oxygen species (ROS) formation, membrane disruption, and pro-inflammatory cytokine release (Caillon et al., 2016; Masters et al., 2010; Westwell-Roper et al., 2014), we analyzed gene expression in INS-1E β-cells to identify key affected pathways. Consistent with previous reports implicating hIAPP in β-cell dysfunction and oxidative stress (Back and Kaufman, 2012; Casas et al., 2007; Rodríguez-Comas et al., 2020), treatment with exogenous hIAPP led to downregulation of *Slc2a2* and *Hspa5* as well as upregulation of *IL-1β*, indicating impaired glucose metabolism, ER stress, and enhanced inflammatory signaling. Notably, QBP1 co-treatment reversed these transcriptional changes by restoring *Slc2a2* and *Hspa5* expression and reducing *IL-1β* levels, thereby mitigating hIAPP-induced stress. QBP1 also upregulated *Pdx1*, a key regulator of β-cell identity and insulin gene transcription, further supporting its protective role. These effects align well with previous studies highlighting the therapeutic value of chaperone-based interventions in preserving β-cell homeostasis (Cadavez et al., 2014; Guo et al., 2013; Montane et al., 2016; Polonsky, 2009). In particular, *Hspa5* induction by QBP1 likely contributes to ER stress relief by enhancing protein folding capacity. Altogether, these findings suggest that QBP1 enhances β-cell resilience by modulating stress, inflammatory, and metabolic gene networks disrupted by hIAPP aggregation. Combined with its positive impact on cell viability, these molecular effects position QBP1 as a promising candidate for counteracting hIAPP-induced β-cell dysfunction.

An important consideration that emerges from our findings is the distinction between intracellular and extracellular hIAPP-induced toxicity (Raleigh et al., 2017). While QBP1 provided protection in both contexts—likely by preventing or reducing oligomer formation—the underlying mechanisms may differ significantly. Intracellularly, hIAPP aggregates form within the secretory pathway or the cytosol, potentially disrupting organelle function and intracellular signaling, including interference with K-ATP channel activity and impaired insulin secretion (Haataja et al., 2008; Kiriyama and Nochi, 2018; Soty et al., 2011). Extracellularly, aggregates may interact with the plasma membrane or are internalized, potentially triggering membrane destabilization, ER stress signaling, and reduced cell viability, even impacting pathways such as mTOR (Casas et al., 2007; Kanatsuka et al., 2018). These distinct modes of toxicity suggest that hIAPP acts through multiple, localization-dependent mechanisms. Although both ultimately converge on β-cell dysfunction, the initial events remain incompletely understood. Importantly, our data show that QBP1 is effective in both models, underscoring its therapeutic versatility. Its dual protective effect likely stems from its capacity to block the formation of toxic oligomers, regardless of their origin. Elucidating these context-specific pathways in the future will be key to refining QBP1-based strategies and tailoring interventions to different stages of the islet amyloid pathology.

Finally, we have investigated the molecular interaction between QBP1 and hIAPP using MD simulations (Case et al., 2020; De Vries et al., 2010; Montane et al., 2017). While QBP1 is known for its broad specificity—effectively inhibiting amyloid formation in several proteins, including TDP-43 (where it primarily targets Q/N-rich segments, (Mompeán et al., 2019)), its inhibition of hIAPP aggregation appears to involve additional molecular determinants. Given hIAPP’s distinct amyloidogenic regions and hydrophobic core (Bakou et al., 2017; Fortier et al., 2022), we hypothesized that the specific sequence and conformation of QBP1 enable it to disrupt β-sheet formation by engaging critical hydrophobic residues, thereby preventing fibril assembly. Our simulations and binding energy analyses provided compelling evidence that QBP1 binds to hIAPP with high affinity and stability, effectively targeting key amyloidogenic segments (residues 8–20, 20–29, and 30–37) essential for β-sheet formation (Nilsson and Raleigh, 1999; Scrocchi et al., 2003). This inhibitory effect is closely linked to the structural conformation of QBP1—characterized as a β-strand-like element followed by a turn featuring a *trans* proline, and stabilized by hydrophobic interactions among nonpolar residues (Mompeán et al., 2019; Ramos-Martín et al., 2014). The prominent role of van der Waals interactions, particularly through W and F residues, underscores the importance of hydrophobic forces in QBP1-mediated amyloid inhibition. Moreover, the peptide’s well-aligned binding conformation optimizes π-H bonding, further reinforcing its inhibitory effect on amylin aggregation.

It should be noted that the absolute value of the predicted binding free energy of QBP1 (–5.70 kcal/mol) is relatively modest compared to those of high-affinity drug candidates, which typically exhibit ΔG_bind_ values below –8.0 kcal/mol (Copeland, 2013; Pantsar and Poso, 2018). However, this benchmark is primarily applicable to small-molecule inhibitors targeting well-defined binding pockets (*e.g.*, kinase inhibitors or enzyme active-site binders). In the context of peptide-based inhibitors like QBP1—designed to interfere with dynamic and transient protein aggregation interfaces—such high affinities are not always necessary to achieve therapeutic efficacy. As shown in Table 1 (*see* the Results section), QBP1 exhibits a substantial enthalpic contribution to binding, largely driven by van der Waals interactions and hydrophobic contacts. This favorable interaction energy is partially counteracted by a significant entropic penalty, likely arising from the intrinsic flexibility of short peptides and solvent fluctuations—factors that commonly introduce variability in computed ΔG_bind_ values. These entropic effects, coupled with methodological approximations inherent to molecular dynamics and free energy calculations, can lead to underestimation or overestimation of the total binding energy. In addition, the ΔG_bind_ that we have reported reflects the most probable binding mode sampled in our simulations but does not preclude the existence of alternative, less populated conformations with potentially stronger affinities. Despite its moderate ΔG_bind_, QBP1 consistently targets amyloidogenic regions of hIAPP with a stable conformation that optimizes π-H and hydrophobic contacts. This supports its *in vitro* activity and highlights its robust inhibitory mechanism.

By contrast, SC-M8 and SC-M1 exhibited weaker and less probable interactions with hIAPP, reflecting suboptimal binding and reduced inhibitory capacity (Okamoto et al., 2009; Ramos-Martín et al., 2014). Their lack of defined structure and hydrophobic core (Ramos-Martín et al., 2014) likely limits their ability to effectively disrupt aggregation. Consistently, ThT aggregation kinetics showed that SC peptides delay amyloid formation, but only partially, mirroring previous findings in TDP-43 models where inhibition was also incomplete (Mompeán et al., 2019). These results reinforce that sequence specificity and precise conformational alignment are critical for the superior inhibitory function of QBP1. Noteworthy, light microscopy revealed distinct aggregation patterns under QBP1 and SC treatment. QBP1 reduced large, amorphous hIAPP aggregates (Azzam et al., 2018; Niu et al., 2020; Pytowski et al., 2020)—suggesting effective fibril disruption—while SC peptides promoted spherical or ring-like aggregates and droplet-like features consistent with LLPS-derived condensates (Pytowski et al., 2021, 2020). Although these SC-induced structures were ThT-positive, they lacked A11 and OC reactivity, suggesting non-canonical, antibody-inaccessible conformations. Combined with their structural features and solvation profiles, this suggest that in these cases the amyloidogenic pathway is not fully completed, instead favoring LLPS or alternative compact assemblies (Ramos et al., 2023; Vendruscolo and Fuxreiter, 2022). Overall, these findings show that SC peptides not only delay fibril formation but also reshape the aggregation pathway, highlighting their unique structural influence on hIAPP assembly.

Despite their mechanistic interest, SC variants fall short of QBP1 in terms of stability, specificity, and inhibitory potency. QBP1 consistently showed robust anti-amyloidogenic activity at significantly lower concentrations than other reported peptides (Wirth et al., 2023; Xu et al., 2022). For instance, FLPNF required around 200 µM to reduce hIAPP-induced cytotoxicity (Shi et al., 2019), while a only 10 μM of QBP1 was needed. Similarly, D-ANFLVH needed a 10-fold molar excess relative to IAPP to prevent toxicity in Rin1056A β cells (Potter et al., 2009; Wijesekara et al., 2015), whereas equimolar QBP1 was sufficient in our model. These findings suggest that QBP1 could be one of the most potent peptide inhibitors of hIAPP aggregation and toxicity reported so far. Reducing the required dose is key when designing new peptide treatments as it significantly reduces negative side effects and production costs.

Remarkably, QBP1 has also demonstrated efficacy in unrelated aggregation-prone systems, such as polyQ-driven toxicity in COS-7 cells, where *Antp*-QBP1 reduced inclusion bodies formation at concentrations as low as 20 μM (Popiel et al., 2007). Originally designed to target expanded polyQ stretches, QBP1 achieved near-complete *in vitro* inhibition at a 3:1 molar ratio with thio-Q62 (thio-Q62: QBP1) (Popiel et al., 2011); though higher doses were needed for more aggregation-prone targets like thio-Q81 (1:10, thio-Q81: QBP1) (Nagai et al., 2000), presumably due to the higher aggregation propensity and increased number of accessible binding sites of thio-Q81. In light of these concentration-dependent effects, the next crucial step is to assess the therapeutic potential of PTD-QBP1 in a mammalian T2D model—which may require modifications to enhance its stability, half-life, and *in vivo* inhibitory effectiveness. This is particularly relevant given the limitations of current T2D treatments and underscores the need for alternative strategies to prevent and delay disease progression by preserving β-cell function. Finally, based on the existing sequence similarity between hIAPP and Aβ, and the evidence of cross-seeding between them (Krotee et al., 2018; Miklossy et al., 2010; Pruzin et al., 2018), QBP1 may also inhibit the cross seeding of Aβ with hIAPP, which would open new avenues for its application to T2D-associated Alzheimer’s disease—a hypothesis we aim to explore in future studies.

## Conclusions

Our results stablish QBP1 as a potent and versatile conformational inhibitor of amyloidogenesis, demonstrating comparable or superior efficiency for human IAPP compared to other target amyloids and other inhibitors. By effectively preventing amyloidogenesis from its outset, QBP1 not only limits the formation of toxic species but also alleviates the harmful impact of human IAPP on pancreatic cells. Consequently, QBP1 emerges as a promising therapeutic candidate for the prevention and management of type 2 diabetes, offering new hope in the search for effective treatments targeting this and other amyloid-related pathologies. Furthermore, the proposed molecular mechanism involved in this interaction paves the way for its future improvement by rational design, if needed.

## Acknowledgments

This work was supported by grants from the Spanish Ministry of Science and Innovation (MICINN: PID2020-117847RB-I00/AEI/10.13039/501100011033 and the Spanish Research Council (CSIC: PIE 202420E226) to MC-V. The authors also thank Prof. Javier DeFelipe, Dr. José Rodrigo Rodríguez Sánchez, Nicolás Cano Astorga, and Sergio Plaza Alonso from CSIC for their outstanding technical assistance with TEM imaging. We are particularly grateful to Dr. Douglas Vinson Laurents (CSIC) for his valuable scientific input and insightful comments during the preparation of the manuscript.

