## Supplementary Information 1 for "A potential anti-amyloidogenic therapy for type 2 diabetes based on the QBP1 peptide"

**Table S1:**

| **Gene** | **Species** | **Fw** | **Rv** |
| --- | --- | --- | --- |
| ***Pdx1*** | Rattus norvegicus | AACTGTCAAAGCGATCTGGG | GCGTAGTACTGCTCCTCACT |
| ***IL-1β*** | Rattus norvegicus | ACAGCAATGGTCGGGACATA | CTGAGAGACCTGACTTGGCA |
| ***Slc2a2*** | Rattus norvegicus | GATGCTACACTTGGCTCAGC | CACAAGCAGCACAGAGACAG |
| ***Hspa5*** | Rattus norvegicus | CATCTGGGTGGGGAAGACTT | AATCCTCGCTTGATGCTGAG |
| ***Rn18s*** | Rattus norvegicus | CGTTCTTAGTTGGTGGAGCG | CCGGACATCTAAGGGCATCA |

***Table S1.* Oligonucleotide primers used for gene expression analysis by RQ-PCR.** Summary of the forward (Fw) and reverse (Rv) primers used for real-time quantitative PCR (RQ-PCR) analysis of *Pdx1*, *IL-1β*, *Slc2a2*, *Hspa5*, and *Rn18s* (*18S rRNA*, reference gene) in *Rattus norvegicus*. The sequences were designed to ensure optimal specificity and amplification efficiency in gene expression studies.

**Figure S1:**

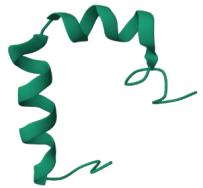

***Figure S1.* Three-dimensional structure of the amylin conformer (PDB ID: 2L86) used in molecular binding simulations.** Structural representation of human amylin (hIAPP) based on NMR solution structure (PDB ID: 2L86) (Nanga et al., 2011). This conformer was used as the reference model in molecular dynamics simulations to study its interactions with QBP1 and scrambled variants.

**Table S2:**

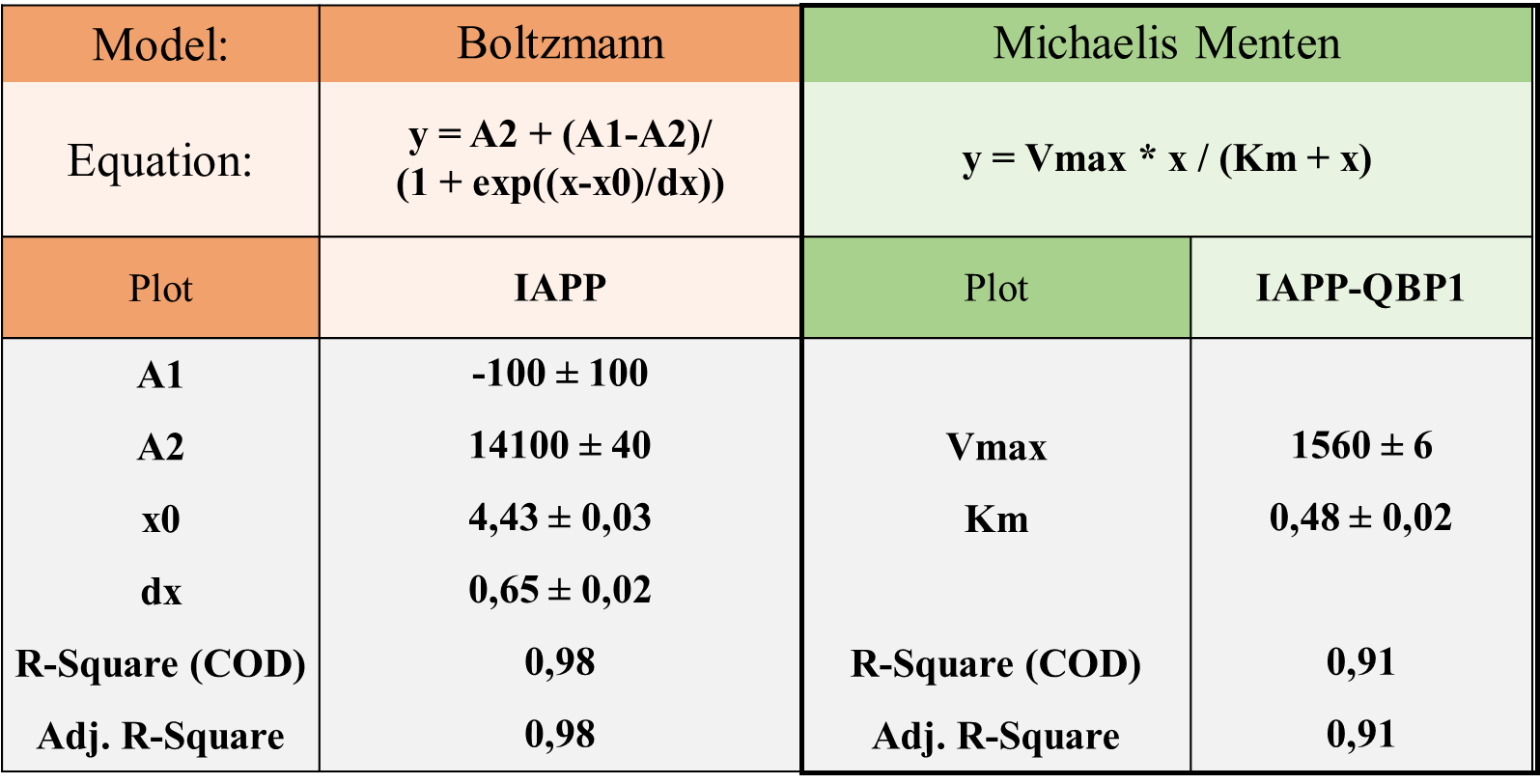

***Table S2.* Kinetic modelling of human IAPP aggregation in the absence and presence of QBP1.** Fluorescence intensity data were fitted to non-linear models to compare the aggregation dynamics of human IAPP alone (Boltzmann model) and IAPP-QBP1 (Michaelis-Menten model). The Boltzmann fit for IAPP alone yielded a maximum fluorescence intensity (A2) of 14100 a.u., reflecting extensive amyloid formation. In contrast, the Michaelis-Menten fit for IAPP-QBP1 showed a drastic reduction in maximum fluorescence intensity (V_max_ = 1560 a.u.), indicating strong inhibition of amyloid polymerization. Additionally, the aggregation half-time (t₁_/_₂) was obtained by solving the Boltzmann and Michaelis-Menten equations, revealing a reduction from 4.4 h (IAPP alone) to 0.49 h (QBP1). All kinetic modelling and curve fitting were performed using OriginPro software.

**Table S3:**

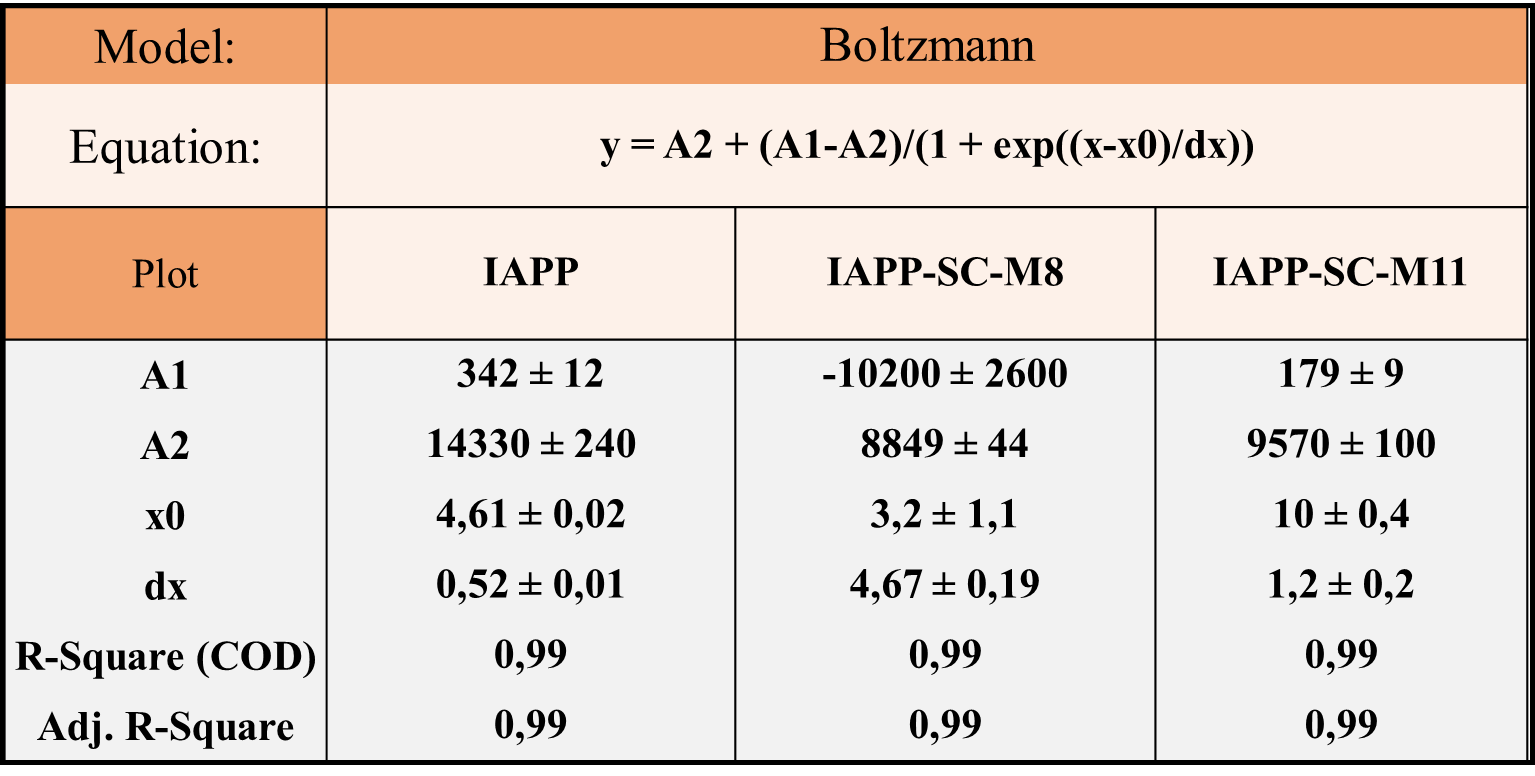

***Table S3.* Kinetic modelling of human IAPP aggregation in the presence and absence of scrambled variants (SC-M8 and SC-M11).** Fluorescence intensity data were fitted to a Boltzmann model, showing that SC-M8 and SC-M11 reduced amyloid formation but were less effective than QBP1. The maximum fluorescence intensity (A2) decreased from 14330 a.u. (IAPP alone) to 8849 a.u. (SC-M8) and 9570 a.u. (SC-M11). Additionally, the aggregation half-time (t_₁/₂_) was obtained by solving the Boltzmann equation, revealing an increase from 4.61 h (IAPP alone) to 8.62 h (SC-M8) and 9.91 h (SC-M11). All kinetic modelling and curve fitting were performed using OriginPro software.

**Figure S2:**

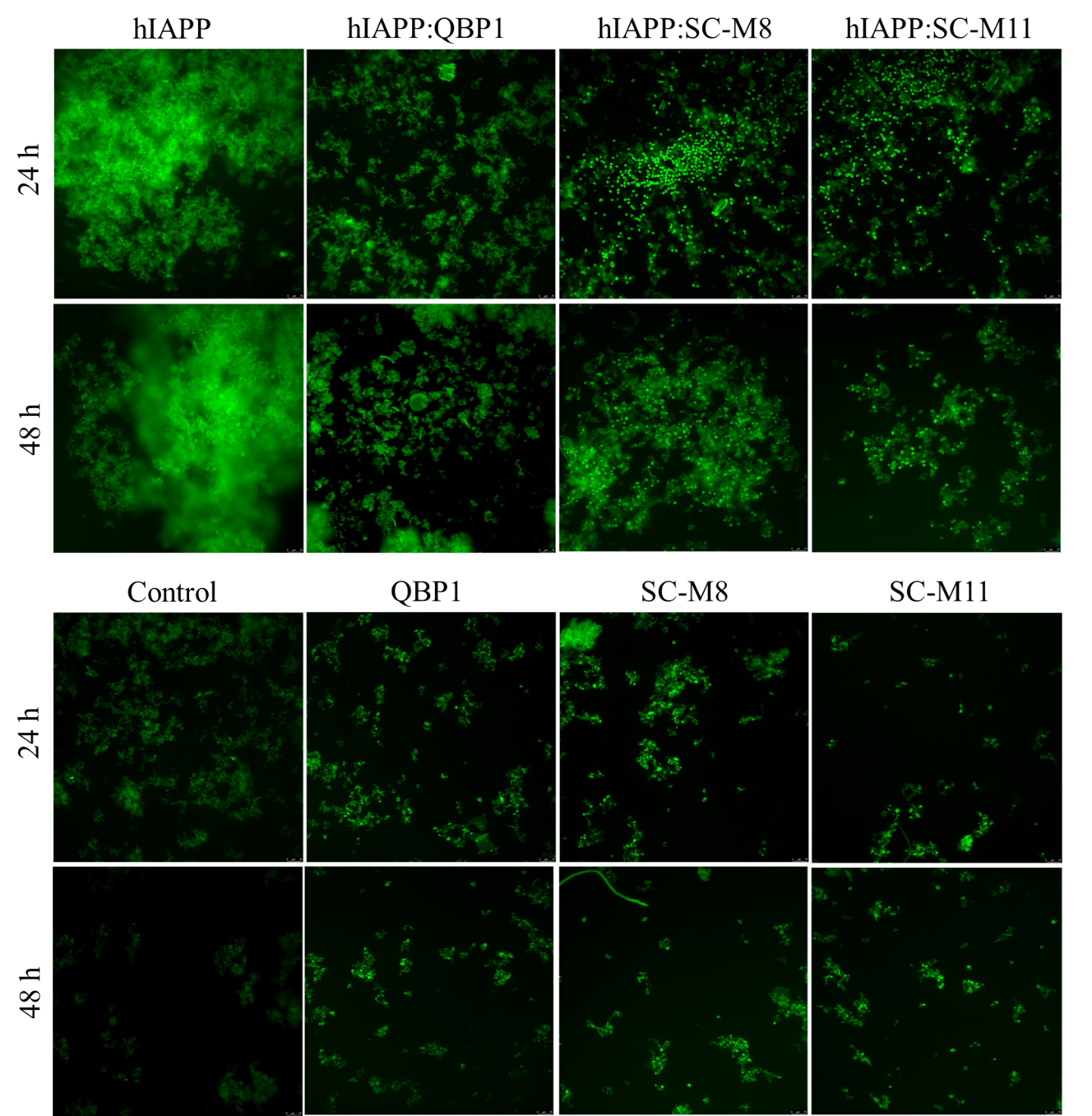

***Figure S2.* Fluorescence microscopy of ThT-stained hIAPP samples at 24 and 48 hours, with or without inhibitory peptides (QBP1, SC-M8 and SC-M11).** Fluorescence microscopy images (10× magnification) of ThT-stained samples reveal amyloid aggregation profiles under different experimental conditions. hIAPP alone formed abundant, dense fluorescent deposits consistent with extensive fibril formation. Co-incubation with QBP1 led to a marked reduction in fluorescence, suggesting potent inhibition of aggregation. Samples treated with SC peptides (SC-M8 and SC-M11) displayed intermediate fluorescence intensity, suggesting partial suppression of amyloid formation. Control conditions—including buffer alone, QBP1 alone, and SC peptides alone—exhibited negligible ThT fluorescence at both time points. This confirms that the observed signals in the treated samples are specifically dependent on hIAPP aggregation and/or its interaction with the respective inhibitors. Scale bars: 75 μm. Representative images from three independent experiments.

**Figure S3:**

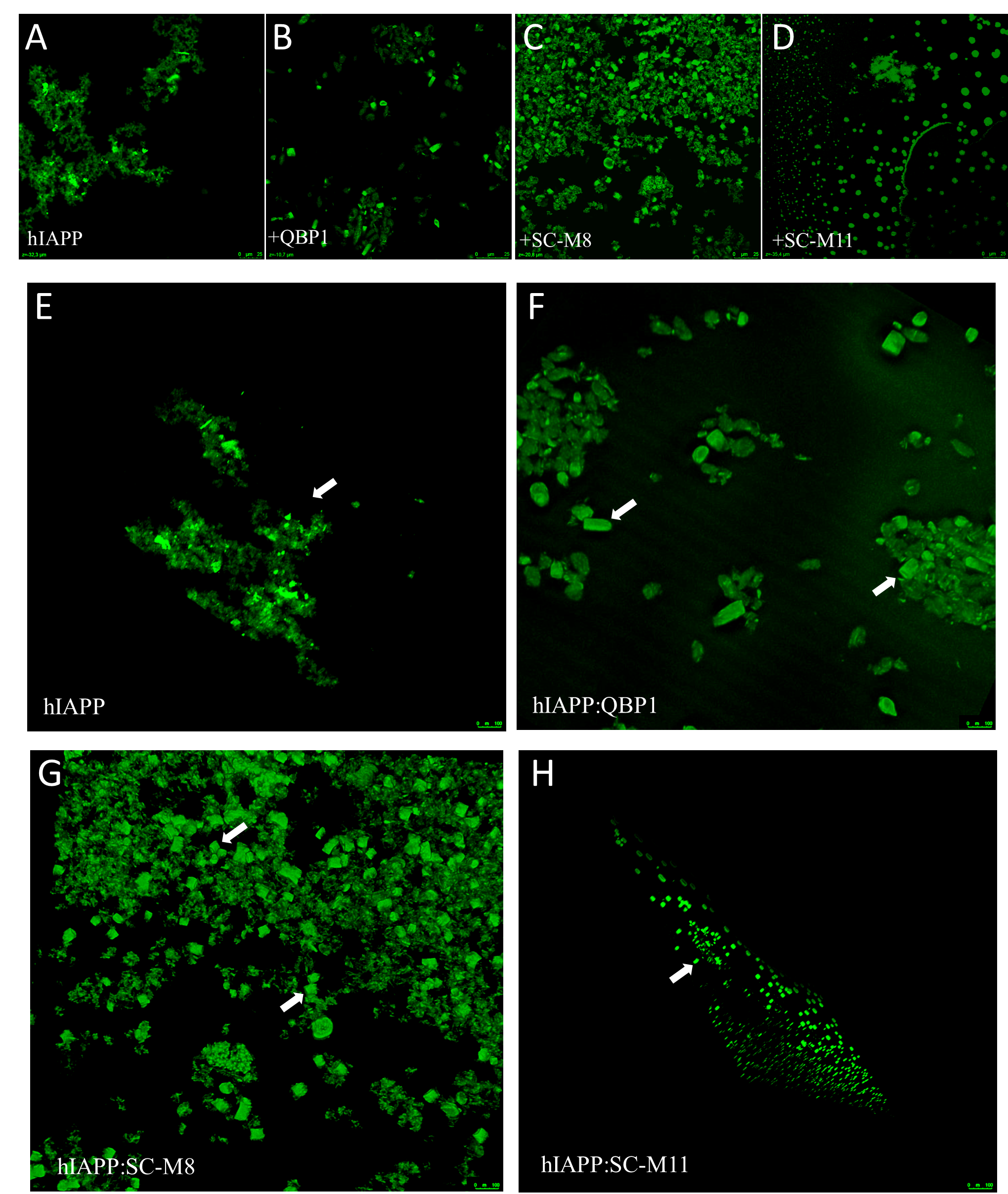

***Figure S3.* Confocal fluorescence microscopy and 3D reconstructions of hIAPP aggregates under different conditions.** Images were acquired using a Leica Stellaris 8 STED super-resolution microscope with a 40× oil-immersion objective after 7 days of incubation. Panels A–D display 2D confocal fluorescence images of ThT-stained hIAPP aggregates, while panels E–H show the corresponding 3D reconstructions generated from confocal z-stacks, providing a more detailed spatial visualization of aggregate morphology (highlighted by white arrows). **(A, E)** hIAPP alone formed dense, amorphous ThT-positive aggregates, consistent with extensive fibril formation. **(B, F)** The hIAPP: QBP1 condition showed fragmented and less organized structures, suggesting that QBP1 disrupts fibril elongation. **(C, G)** hIAPP:SC-M8 samples induced the formation of compact, well-defined aggregates, distinct from the large amorphous clusters seen in hIAPP alone. **(D, H)** hIAPP:SC-M11 formed predominantly spherical or ring-like structures; 3D reconstructions revealed hollow, droplet-like morphologies, indicative of a phase separation mechanism distinct from classical fibrillogenesis. White arrows indicate representative structures of interest in each condition. These results support the notion that QBP1 and SC peptide variants modulate hIAPP aggregation, redirecting it toward structurally distinct endpoints. Scale bars: 25 μm. Representative images from three independent experiments.

**Figure S4:**

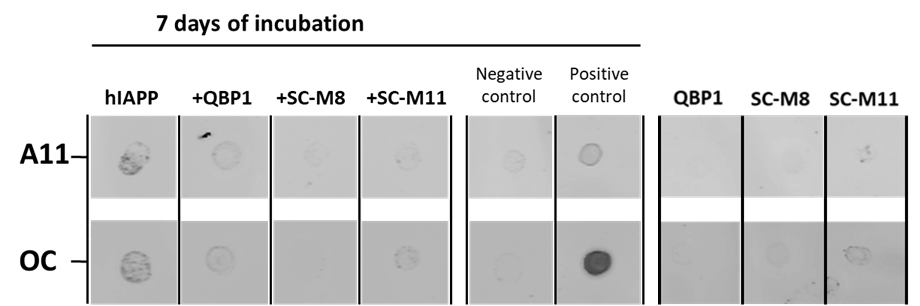

***Figure S4.* Immunodot-Blot analysis of hIAPP aggregation in the presence of QBP1 and scrambled variants.** Dot-blot analysis of hIAPP (70 μM) alone or co-incubated with QBP1, SC-M8, or SC-M11 (1:5 molar ratio) after 7 days of incubation, using A11 (oligomer-specific) and OC (fibril-specific) antibodies. hIAPP alone showed strong A11 and OC signals, confirming the presence of oligomeric and fibrillar species. In contrast, QBP1, SC-M8, and SC-M11-treated samples exhibited markedly reduced immunoreactivity, indicating inhibition of amyloid progression. As expected, control reactions containing QBP1, SC-M8, or SC-M11 alone (rightmost panels) showed negligible A11 or OC signals, confirming that these peptides do not spontaneously aggregate under the experimental conditions. Controls: BSA (non-amyloidogenic, negative) and Aβ42 (amyloidogenic, positive). Representative images from three independent experiments.

**Figure S5:**

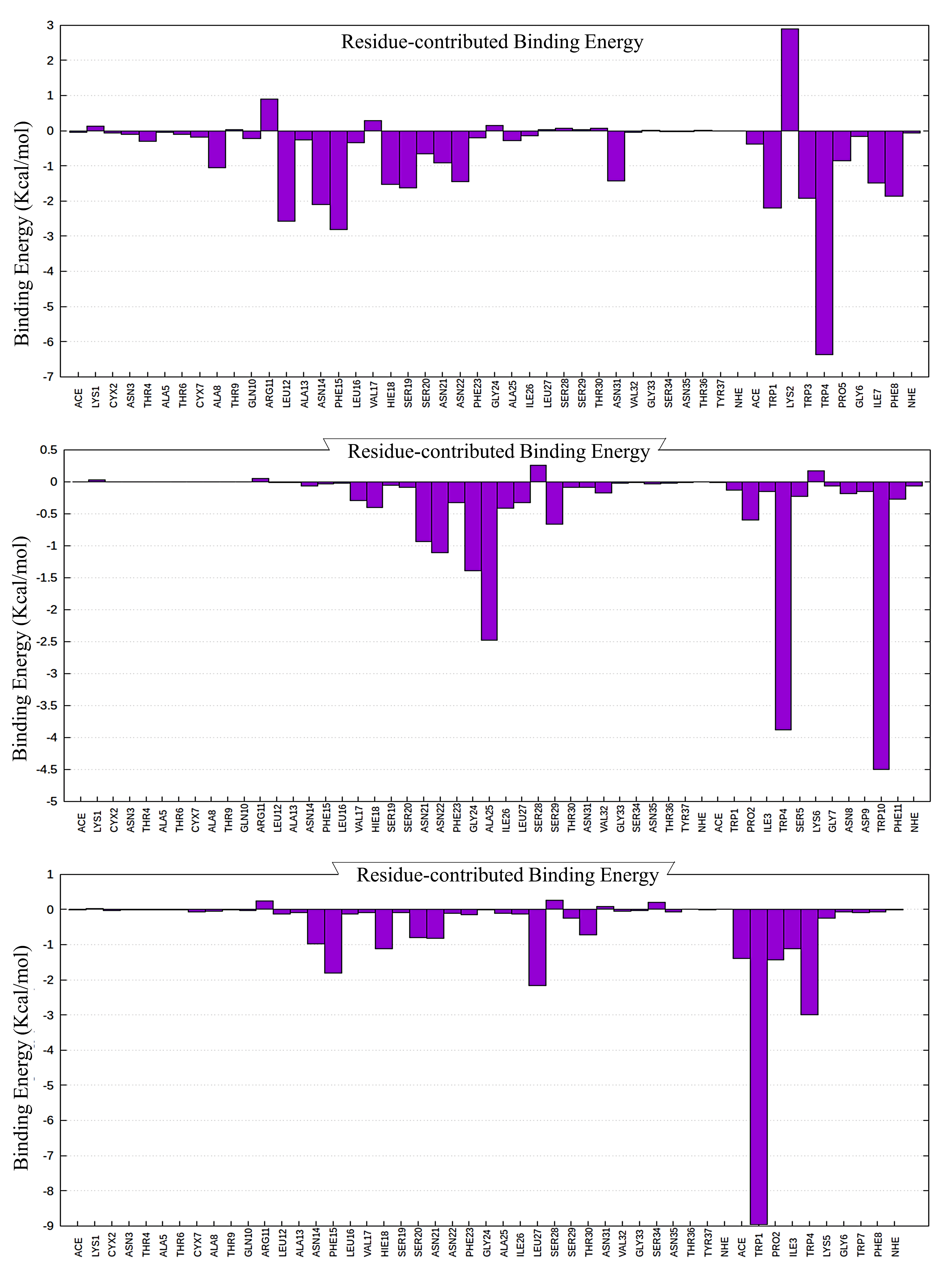

***Figure S5.*** ***Per*-residue contribution to the binding free energy for the amylin-QBP1, amylin-SC-M11, and amylin-SC-M8 complexes.** *Per*-residue decomposition of binding free energy for the amylin-QBP1 (top panel), amylin-SC-M11 (middle panel), and amylin-SC-M8 (bottom panel) complexes. The initial residues correspond to amylin, while the final residues represent QBP1, SC-M11, or SC-M8. Negative values indicate stabilizing interactions, whereas positive values suggest destabilizing effects.

**Figure S6:**

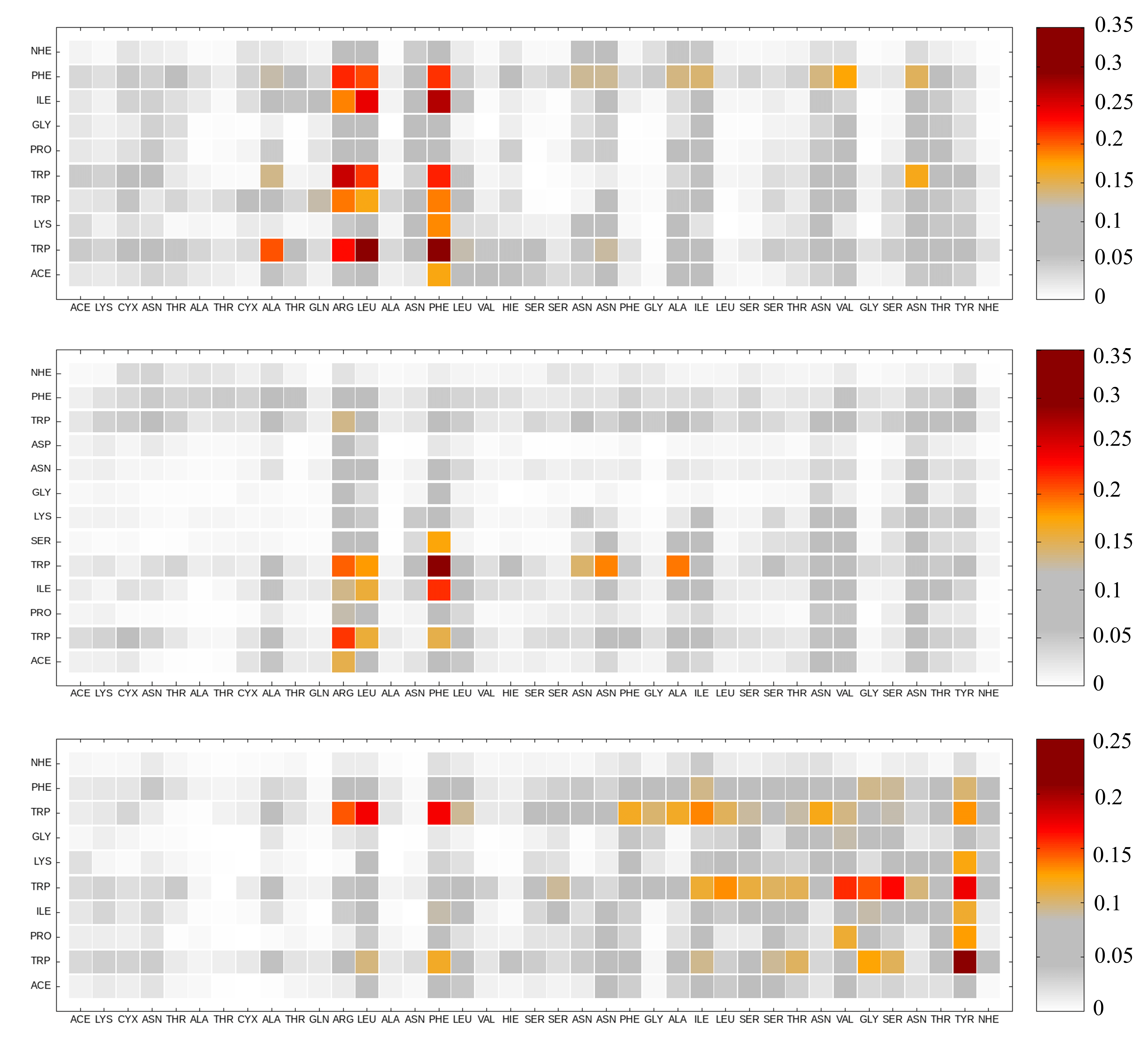

***Figure S6.* Averaged contact maps for amylin-QBP1, amylin-SC-M11), and amylin-SC-M8 complexes.** Averaged contact maps derived from molecular dynamics simulations showing the most probable molecular interactions in the amylin-QBP1 (top panel), amylin-SC-M11 (middle panel), and amylin-SC-M8 (bottom panel) complexes. The color scale represents the probability of contact formation, with red indicating high probability and grey indicating low probability. These maps highlight key binding regions and differences in interaction patterns, reinforcing the consistency of the identified binding sites across simulations.

**Figure S7:**

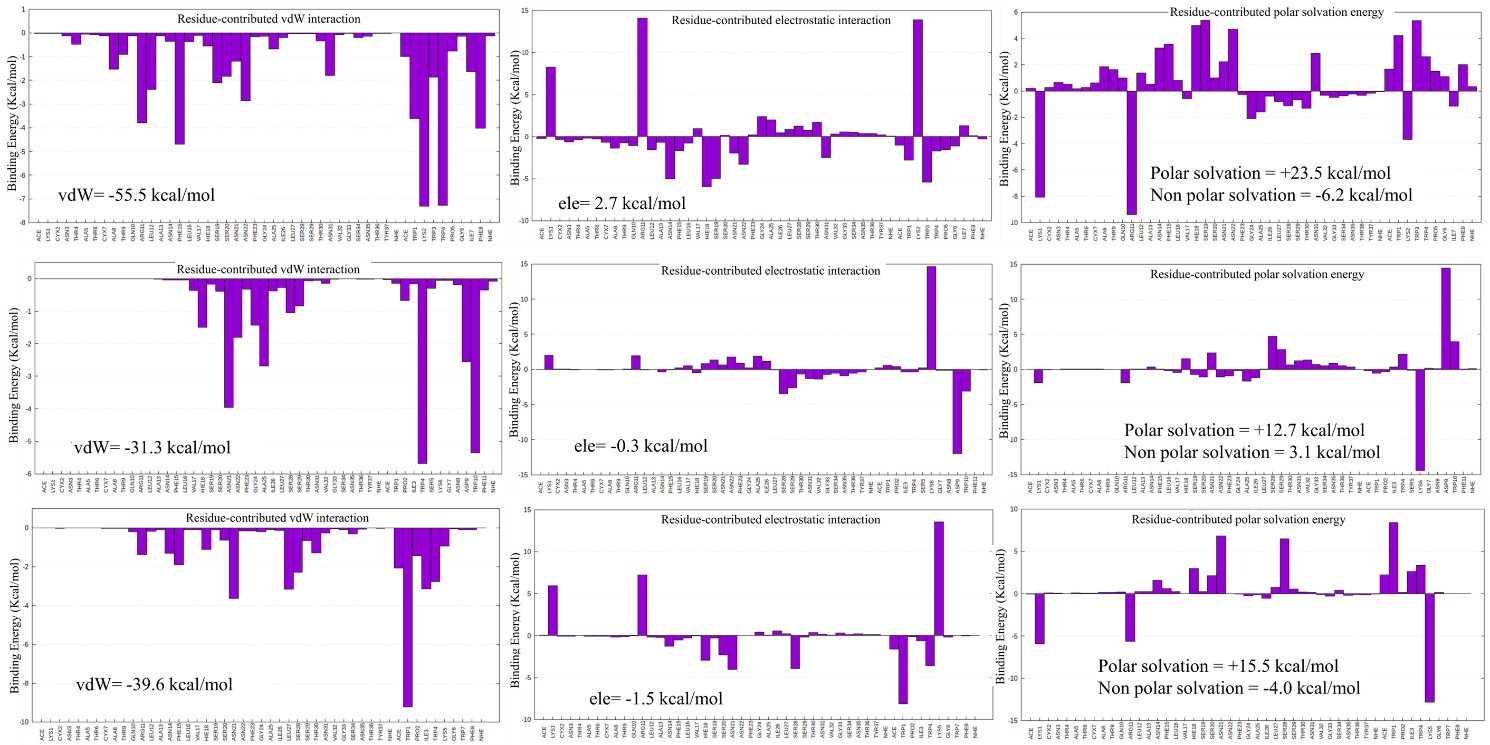

***Figure S7.* Per-Residue decomposition of binding energy components in amylin-QBP1, amylin-SC-M11, and amylin-SC-M8 Complexes.** *Per*-residue decomposition of van der Waals interactions (left panels), electrostatic interactions (middle panels), and polar solvation energy (right panels) for the amylin-QBP1 (top), amylin-SC-M11 (middle), and amylin-SC-M8 (bottom) complexes. The initial residues correspond to amylin, while the final residues represent QBP1, SC-M11, or SC-M8. QBP1 exhibits the strongest van der Waals contributions, particularly in hydrophobic and aromatic residues, enhancing its binding stability. In contrast, polar solvation energy is consistently unfavorable across all complexes, with SC variants showing greater destabilization. The abbreviations ACE and NHE denote the N-terminal acetyl and C-terminal amide blocking groups, respectively.

**Figure S8:**

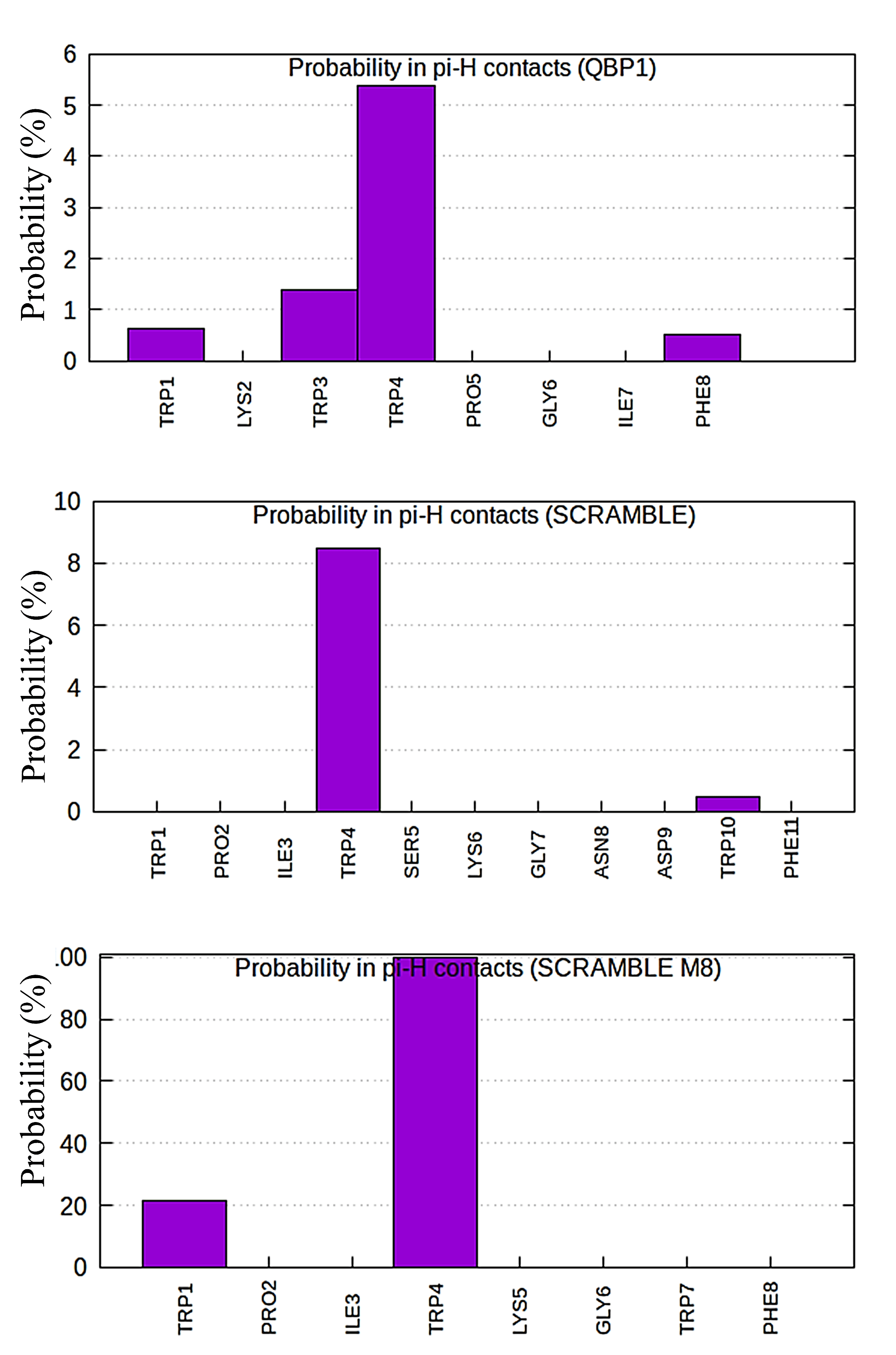

***Figure S8*. Probability of π-H bond formation in amylin-QBP1, amylin-SC-M11, and amylin-SC-M8 complexes.** Bar plots showing the probability of π-H bond formation for individual residues in the amylin-QBP1 (top panel), amylin-SC-M11 (middle panel), and amylin-SC-M8 (bottom panel) complexes. In the y-axis we represent the probability (%), while in the x-axis the participating residues. QBP1 exhibits a higher occurrence of π-H interactions, particularly at W and F residues, compared to SC-M11 and SC-M8, contributing to its superior stability and inhibitory efficiency against amylin aggregation.
